# ERSAtool: A User-Friendly R/Shiny Comprehensive Transcriptomic Analysis Interface Suitable for Education

**DOI:** 10.1101/2025.06.20.660710

**Authors:** Sujith Taridalu, Ayyappa Kumar Sista Kameshwar, Masako Suzuki

## Abstract

RNA sequencing (RNA-seq) has become an essential technology for assessing gene expression profiles in biomedical research. However, the coding complexity of RNA-seq data analysis remains a significant barrier for students and researchers without extensive bioinformatics expertise. We present the <u>**E**</u>ducational <u>**R**</u>NA-<u>**S**</u>eq <u>**A**</u>nalysis <u>**tool**</u> (ERSAtool), a comprehensive R/Shiny interface that provides an intuitive graphical visualization of the complete RNA-seq analysis workflow. The application is built on established Bioconductor packages and upholds high standards in analyses while significantly reducing the technical expertise required to conduct sophisticated transcriptomic analyses. ERSAtool supports various input formats, such as raw count matrices and STAR alignment outputs. It generates sample information metadata through direct integration with the Gene Expression Omnibus (GEO) provided by the National Center for Biotechnology Information (NCBI). The application guides users through normalization, data visualization, differential expression analysis, and functional interpretation using Gene Ontology (GO) and Gene Set Enrichment Analysis (GSEA). All results can be compiled into comprehensive, downloadable reports that enhance reproducibility and knowledge sharing. The design includes targeted features that facilitate educational use, making it especially useful for teaching transcriptomics in undergraduate to graduate-level bioinformatics courses. By connecting command-line bioinformatics tools with accessible graphical interfaces, ERSAtool improves accessibility to advanced transcriptomic analysis capabilities, potentially accelerating discoveries across various biological fields. The source code for the ERSAtool package is available at https://github.com/SuzukiLabTAMU/ERSAtool and is released under the GNU General Public License v3.0 (GPL-3.0).

## 1. Introduction

In current biomedical research, RNA-seq is the gold standard for transcriptome analysis, facilitating comprehensive profiling of gene expression across various biological systems (Conesa et al., 2016; Stark et al., 2019). As sequencing costs decline, RNA-seq technology has become more widely accessible. Yet, the computational analysis of RNA-seq data remains a considerable challenge for many biologists, clinicians, and students who lack specialized expertise in bioinformatics and statistics (Conesa et al. 2016). Although the Bioconductor project offers powerful open-source R packages for RNA-seq analysis, its command-line interface poses a significant learning barrier for researchers without programming experience (Huber et al. 2015; Love et al. 2014).

Several RNA-seq pipelines have attempted to address this gap. Still, limitations persist due to factors such as restricted analytical scope, limited customizable options, data privacy concerns related to external web servers, and complex installation requirements (Gu et al., 2016; Nelson et al., 2017; Russo et al., 2016). We developed a comprehensive and interactive Shiny application, the <u>**E**</u>ducational <u>**R**</u>NA-<u>**S**</u>eq <u>**A**</u>nalysis <u>**tool**</u> (ERSAtool), that supports a complete RNA-seq analysis workflow. It significantly reduces technical barriers and offers a user-friendly interface maintaining the analytical rigor of established Bioconductor packages, including DESeq2 and ClusterProfiler (Love et al., 2014; Wu et al., 2021; Yu et al., 2012).

ERSAtool tackles the frequent challenges that beginners face in RNA-seq analysis, including complex installation steps, creating sample metadata, evaluating and visualizing results, and generating a detailed report that includes code for reproducibility. Additionally, the application supports various input formats, provides direct NCBI GEO integration for metadata management, and guides users through the entire analytical process from normalization to functional interpretation. It enables users to analyze transcriptomic data independently and fosters critical thinking in biological disciplines. We illustrated a workflow for RNA-seq analysis using the ERSAtool in **Figure 1**.

**Figure 1:** RNA-seq analysis workflow using ERSAtool.

## 2. Methods

### 2.1 Implementation and Architecture

ERSAtool is designed as a modular Shiny application in R (≥ 4.1) featuring a component-based architecture that enhances maintainability and facilitates expansion. The application incorporates established Bioconductor packages into a cohesive analytical framework that can be executed through a user-friendly interface.

The core structure is organized around a main entry point (App.R) and global configuration (global.R), with distinct directories for interface components (ui/), server logic (server/), and static assets (www/). User interface components are segregated into module-specific files (e.g., metadata_ui.R, pca_heatmap_ui.R), promoting component reusability and independent development while maintaining a cohesive user experience through the dashboard framework (dashboard.R). Server logic is organized into dedicated four subdirectories: 1) code: implements essential analytical functions for normalization, differential expression, and enrichment analyses; 2) plots: manages visualization rendering for each analysis module; 3) downloads: oversees the export of results and images in formats suitable for publication; and 4) report: enables the creation of detailed analytical reports. This architecture enables independent development of application components and maintains integration across the analytical workflow.

### 2.2 A step-by-step workflow directing users through RNA-seq analysis

#### a) Loading data

ERSAtool requires two data inputs for analysis: sample metadata and raw count data. The sample metadata can be either a user-provided local data file (.txt, .csv, .xlsx) or a Series Matrix File retrieved directly from GEO using the accession number. The first column should contain sample identifiers. Since the application uses DESeq2 (Love et al., 2014) for analysis, the expression data should be in raw count format. The user can upload a row count matrix (.gz, .csv, .txt, .xlsx) or STAR aligner outputs (.out.tab). Both formats should have the first column serving as a gene identifier. For row count matrices, the remaining columns represent sample counts. When the user chooses STAR aligner outputs as count data, it retrieves counts from designated columns (unstranded RNA-seq counts, counts for the first read strand aligned with RNA, or counts for the second read strand aligned with RNA) and extracts sample names from the file names prior to merging them.

#### b) Generating an analysis design formula

After users upload the sample metadata, the interface allows adjustments such as adding, removing, or renaming columns, inline editing of cells, and modifying rows in a paginated table format. Users can also export the modified sample metadata in an Excel format for future use. To generate a design formula, users select the column(s) they wish to include in the analysis.

#### c) Initial Assessment

In the initial phase of the analysis, the application offers a guided assessment utilizing interactive visualizations, including box-whisker plots of raw and normalized counts, a pair plot of principal components (PC1 and PC2), and heatmaps from hierarchical clustering. The boxplot displays variations in sequencing depth across samples before and after normalization, while the PC pair plots and heatmaps reveal similarities in expression profiles among the samples.

#### d) Differential Expression analysis

The application utilizes DESeq2 (Love et al., 2014). Count data undergoes pre-processing validation before being created into a DESeqDataSet. Low-expressed genes are filtered using a minimum threshold of 1 count per sample. The DESeq2 core algorithm then estimates library size factors of each sample, calculates dispersion values, and fits negative binomial generalized linear models for each gene, storing results for visualization and analysis. Differentially expressed genes (DEGs) are identified using the statistical framework of DESeq2. The application includes features with adjustable threshold cutoffs for false discovery rate (FDR)-adjusted p-values and log2-fold changes (logFC), and it accommodates a flexible experimental design for multiple-factor study designs. These accommodations follow established best practices for contrast specification and DEG identification in DESeq2, enabling the analysis of various experimental designs without requiring users to manually create design formulas, contrast matrices or set thresholds for FDR-adjusted p-values and logFC.

#### e) Interpretation of differentially expressed genes

The application supports two types of assays for interpreting gene expression profiles using the *enrichGO* and *gseGO* functions of *clusterProfiler* (Wu et al., 2021; Yu et al., 2012): Gene Ontology (GO) enrichment (GO-enrichment) and GO gene set enrichment analysis (GO-GSEA). GO-enrichment analysis assesses whether the identified significant DEGs are enriched in specific pathways, while GO-GSEA evaluates if a particular gene set is enriched or depleted in the dataset. These results can provide insights into the biological processes or pathways associated with a specific phenotype or condition. The GO-enrichment results are reported separately for the biological process and molecular function categories.

#### f) Report and result file generation

In the final step, the application can compile all analysis results into a comprehensive document that captures the entire analytical process. It also includes a function to export the result files (the list of DEGs and DO terms) in CSV format, as well as the plot images in PNG, JPEG, TIFF, and PDF formats.

To support these capabilities, ERSAtool integrates key R/Bioconductor packages into a unified framework, including core functionality packages (*shiny, shinydashboard, rmarkdown*) (Chang et al., 2024), data processing tools (*data.table, dplyr, readr, readxl*) (Barrett et al., 2024; Wickham & Bryan, 2024; Wickham et al., 2024a, 2024b), RNA-seq analysis packages (*DESeq2, GEOquery)* (Davis & Meltzer, 2007; Love et al., 2014), visualization libraries (*ggplot2, plotly, pheatmap, PCAtools*) (Kolde, 2024; Marini & Binder, 2019), and functional analysis tools (*clusterProfiler, enrichplot, fgsea*) (Sergushichev, 2017; Wu et al., 2021; Yu, 2018; Yu et al., 2012). ERSAtool requires R (version 4.1 or later), RStudio (recommended), and an internet connection for initial package installation and retrieval of GEO data. The application is launched by executing App R through the RStudio interface or via shiny::runApp (Chang et al., 2024). Upon initial execution, the application automatically detects missing package dependencies. It then downloads and manages all dependencies through runtime package verification, thereby reducing installation barriers for users with varying coding skills. The application is designed for local deployment, facilitating analysis of sensitive data without external transfer while maintaining accessibility for users with basic R installation knowledge. It is also suitable for educational settings with limited computational resources. A detailed step-by-step user guide is supplemented as **Supporting Material 1**.

### 2.3 Educational functions and tools

ERSAtool is designed to guide students and researchers through a structured RNA-seq workflow, featuring dynamic visualizations that update as data is inputted. Dedicated interpretation text boxes beneath each visualization prompt users to document their insights during the analysis process. The entire workflow, including all user interpretations, can be exported as a static HTML report suitable for documentation or collaborative sharing, creating an efficient yet flexible approach that combines bioinformatics analysis with guided critical interpretation of biological significance. This structured framework enables users to concentrate on biological meaning while the tool manages computational complexity. In addition, the function to export the DEG or enriched pathway results allows the users to perform further analysis using web-based analyses, such as EnrichR (Chen et al., 2013), STRING (Szklarczyk et al., 2023), GeneVenn (Pirooznia et al., 2007), and more. For advanced learners, the application displays all the code used at every stage, assisting them in writing their own code later.

### 3.0. Use Cases

#### Case Study: Identifying changes in tissue-specific gene expression of adipose precursor cells resulting from high-fat diet intake

We used a subset of a publicly available dataset (GSE273569 (Wing et al., 2025)) for this case study which focuses on male mice. The original study tested the obesogenic effects of dietary oleic acid on adipocyte precursor cells (APCs). The authors performed transcriptional analysis on APCs from subcutaneous white adipose tissue (SWAT) and visceral white adipose tissue (VWAT), which were sorted from mice fed a standard diet or a 60% kcal lard high-fat diet (HFD) for 3 days, a period when APC proliferation peaks. The authors found that HFD feeding increases VWAT APC proliferation in males (Wing et al., 2025). While the authors reported their findings with a sophisticated analysis, we performed a straightforward analysis focusing on male samples as a case study. We downloaded the unnormalized count and sample matrix files from the Gene Expression Omnibus (GEO). We provided those files as a training dataset (**Supporting Materials 3 and 4**). The primary goal of this analysis was to evaluate the differentially expressed genes in APCs derived from two types of adipose tissue in response to an HFD diet. We created **Figure 2** using the original exported images from the download plot functions. The report HTML file of this analysis has been provided as **Supporting Material 5**.

**Figure 2:** Example RNA-seq analysis on a publicly available dataset using ERSAtool. As a case study, we analyzed a subset of a publicly available RNA-seq dataset (GSE273569) using ERSAtool. In this figure, we show a step-by-step analysis workflow with plots generated at each step. Count distribution analysis of the raw count (**A**) and the normalized count (**B**): While some samples showed slightly lower distributions in raw count, all samples demonstrated comparable count distributions after normalization. Assessing gene expression profile similarity: The similarity of gene expression profiles was evaluated using PCA (**C**) and hierarchical clustering (**D**). Both results indicate that the origin of APC was the primary factor influencing gene expression variations, while diet modulated the gene expression profiles. Differentially expressed gene analysis: among mice fed HFD, we identified 1374 VWAT upregulated and 1291 downregulated genes compared to SWAT (**E**). GO enrichment analysis indicated that these upregulated genes are enriched in cell cycle-related pathways (**F**). The plots included in this figure are the original TIFF files created using the export image function of ERSAtool.

##### a) Count distribution analysis

As a primary assessment of RNA-seq, we examined whether the raw and normalized count distributions were comparable across all libraries (**Figure 2A** and **2B**). Although we noted that some samples exhibited lower raw count distributions (**Figure 2A**), the normalized count distributions were similar among all samples, indicating comparable sequencing depth for the libraries (**Figure 2B**).

##### b) Assessing gene expression profile similarity

We then visualized the gene expression profile similarity between the samples using two independent approaches: principal component analysis (PCA, **Figure 2C**) and hierarchical clustering (**Figure 2D**). Both results showed that the primary factor differentiating the gene expression profiles is the origin of the tissue from which the APCs are derived, with diet being a secondary consideration. The PCA analysis revealed that PC1, which accounts for 66.3% of the variation, dissociates VWAT from SWAT, while PC2, accounting for 18.4% of the variation, differentiates SD from HFD. Although it was grouped within VWAT, one sample from the VWAT-HFD group did not segregate or branch into other VWAT-HFD samples, suggesting that individual variations might exist in response to HFD.

##### c) Differentially expressed gene analysis

We identified differentially expressed genes between VWAT-HFD and SWAT-HFD that exhibit at least 1.5-fold differences with an adjusted p-value of 0.05. Based on this criterion, we have 1374 upregulated and 1291 downregulated genes in VWAT-HFD compared to SWAT-HFD (**Figure 2E**). The upregulated genes are enriched in cell cycle pathways, while the downregulated genes are enriched in immune-response pathways (**Figure 2F**). While upregulation of cell cycle-related pathways in VWAT-HFD is concordant with what the original study reported, this comparison did not account for the gene expression variations between VWAT and SWAT. Therefore, we analyzed VWAT-SD and SWAT-SD and integrated the results to identify DEGs specific to VWAS in response to HFD. In the comparison between VWAT-SD and SWAT-SD, we identified 1281 VWAT upregulated and 1161 downregulated genes compared to SWAT (**Figure 3A**). The lists of tissue origin-dependent DEGs in HFD and SD were exported, respectively. We used GeneVenn, a web-based Venn diagram generator (Pirooznia et al. 2007), to intersect the DEG lists, identifying both overlapping and non-overlapping tissue origin-dependent DEGs. Since we are interested in whether the upregulations in cell cycle-related genes are specific to VWAT, we focused on 408 VWAT-specific upregulated genes for the downstream analysis (**Figure 3B**). We assessed these 408 VWAT-specific upregulated genes using EnrichR, a web-based gene set knowledge discovery tool (Chen et al. 2013), and found enrichments in cell cycle-related pathways among VWAT-specific upregulated genes (**Figure 3C**). This suggests that the upregulation of cell cycle-related pathways in response to HFD is specific to VWAT, aligning with the findings of the original report (Wing et al., 2025). Additionally, our analysis suggested that VWAT-specific upregulated genes are enriched downstream of E2F1 (**Figure 3D**) and enriched Palbociclib (CDK4/6 inhibitor) downregulated genes (**Figure 3E**). The L1000 CRISPR KO analysis results also support that those upregulated genes were downregulated in the DTL, XPO1, HSPA5, CDK6, or CDK4 knock-outs (**Figure 3F**). While the original article does not discuss the enrichment of the CDK4/6-E2F targets in VWAT-specific HFD upregulated genes, CDK4/6-E2F has been implicated in adipogenesis in activating *Pparg* signaling (Abella et al., 2005; Fajas et al., 2002). Our findings indicate that the HFD diet may also differentially affect E2F-CDK4/6 cell cycle regulators based on tissue type, along with LXRα signaling.

**Figure 3:** Example secondary analysis using publicly available web-based tools. We analyzed the results using GeneVenn and EnrichR, which are both publicly available web-based analysis tools. This integrative analysis aims to identify DEGs specific to VWAS in response to HFD. The DEG analysis between the SWAT and VWAT in the SD group identified 1281 VWAT upregulated and 1161 downregulated genes (**A**). The VWAT upregulated and downregulated DEGs in HFD and SD were independently intersected to identify DEGs specific to VWAS in response to HFD using GeneVenn (**B**). Upregulated DEGs specific to VWAS in response to HFD were further analyzed with another web-based gene set enrichment analysis tool, EnrichR (**C-F**). **C**: Reactome Pathways, **D**: TRANSFAC and JASPER Position Weight Matrix, **E**: LINCS L1000 Chemical perturbation consensus signals, **F**: LINCS L1000 CRISPR KO consensus signals.

## 4. Discussion

ERSAtool offers comprehensive learning objectives for analyzing transcriptomic data through a well-crafted, standalone Shiny application. Users develop core analytical skills, including data acquisition and preprocessing by formatting metadata, understanding the structure of RNA-seq analysis, and interpreting their findings. The application teaches normalization principles for effective sample comparisons, employs dimensional reduction techniques for pattern recognition in complex datasets, introduces statistical testing methods for differential expression analysis, and fosters functional interpretation via GO enrichment and GSEA. Conceptually, users gain insights into experimental design, fundamentals of statistics in transcriptomics, the biological interpretation of findings, and strategies for data visualization. The tool also enhances transferable skills such as critical evaluation of results, effective scientific communication, and making well-thought-out analytical decisions. A unique feature that enriches the learning experience is a text box included in each step, which encourages users to interpret their observations during the analysis. These interpretations, along with the conclusion, will be part of the final report, providing support for student assignments and sharing results. In addition, further analysis of exported DEG lists using publicly available tools allows users to perform advanced analyses and interpret the results more deeply. This is conveyed in the case study. By combining computational rigor with guided analytical steps and interpretation prompts, ERSAtool effectively bridges the gap between the complexity of bioinformatics and biological inquiry, empowering users to focus on deriving meaningful insights from gene expression data.

## Acknowledgements

The authors thank Dr. Yoshinori Seki, Ms. Brighton A. Garret, Ms. Kaitlyn Carter, and Mr. William R. Leach for their feedback on the software and manuscript.

## Declarations

### Author Contributions

Conceptualization: MS, Data Curation: ST, AKSK, and MS, Formal Analysis: ST and MS, Investigation: ST and AKSK, Software: ST, Visualization: ST, Supervision: MS, Funding acquisition: MS, Writing—original draft: AKSK and MS, Writing—review and editing: ST, AKSK, and MS.

### Final Approval

All authors have reviewed the manuscript and agree to its submission.

### Ethics Approval

Not applicable.

### Data Availability

All data used in the figures are represented in the Supporting Materials. The source code for the ERSAtool package is available at https://github.com/SuzukiLabTAMU/ERSAtool and is released under the GNU General Public License v3.0 (GPL-3.0).

R Markdown.

Create Elegant Data Visualisations Using the Grammar of Graphics • ggplot2.

Interactive web-based data visualization with R, plotly, and shiny.

## Supporting Information

Supplementary material 1: A step-by-step guide to using ERSAtool

Supplementary material 2: ERSAtool app (**ERSAtool-main.zip**)

Supplementary material 3: Example raw count matrix file (**GSE273569_Males_Unnormalized_Count_Matrix.txt**)

Supplementary material 4: Example sample matrix file (**GSE273569_series_matrix.xlsx**)

Supplementary material 5: Example HTML output (**Tool_06_13_25.html**)

