## Supplementary material for "ERSAtool: A User-Friendly R/Shiny Comprehensive Transcriptomic Analysis Interface Suitable for Education": SupportingMaterial_ERSAtool_16Jun25.docx

Supporting Document 1: A step-by-step guide to using ERSAtool

Supporting Document 2: ERSAtool app (**ERSAtool-main.zip**)

Supporting Document 3: Example raw count matrix file (**GSE273569_Males_Unnormalized_Count_Matrix.txt**)

Supporting Document 4: Example sample matrix file (**GSE273569_series_matrix.xlsx**)

Supporting Document 5: Example HTML output (**Tool_06_13_25.html**)

Supplementary material 1:

### **A step-by-step guide to using ERSAtool**

#### A. Preparation

##### Reading assignment.

Prior to conducting the analysis, users are encouraged to review the following articles to familiarize themselves with RNA-seq technology and DESeq2 analysis.

1. Moderated estimation of fold change and dispersion for RNA-seq data with DESeq2. Love MI et al. Genome Biol. 2014; 15(12) 550 DOI: 10.1186/s13059-014-0550-8, PMID: 25516281
2. Analyzing RNA-seq data with DESeq2. Love MI et al. https://bioconductor.org/packages/devel/bioc/vignettes/DESeq2/inst/doc/DESeq2.html
3. The Hitchhiker’s Guide to RNA Sequencing and Functional Analysis. Chen et al. Brief Bioinform. 2023 Jan 8;24(1):bbac529. doi: 10.1093/bib/bbac529, PMID: 36617463

##### Downloading and installing the app and packages on your computer

1. Install R (<https://www.r-project.org/>) and RStudio (<https://posit.co/download/rstudio-desktop/>)
2. Unzip the supplemented app file (ERSAtool-main.zip) or download the latest ERSAtool app from our GitHub page (recommended, https://github.com/SuzukiLabTAMU/ERSAtool)
3. Open the app.R file with RStudio and start ERSAtool to install all necessary packages by clicking the “Run App” button. This step is only required the first time you use the app.

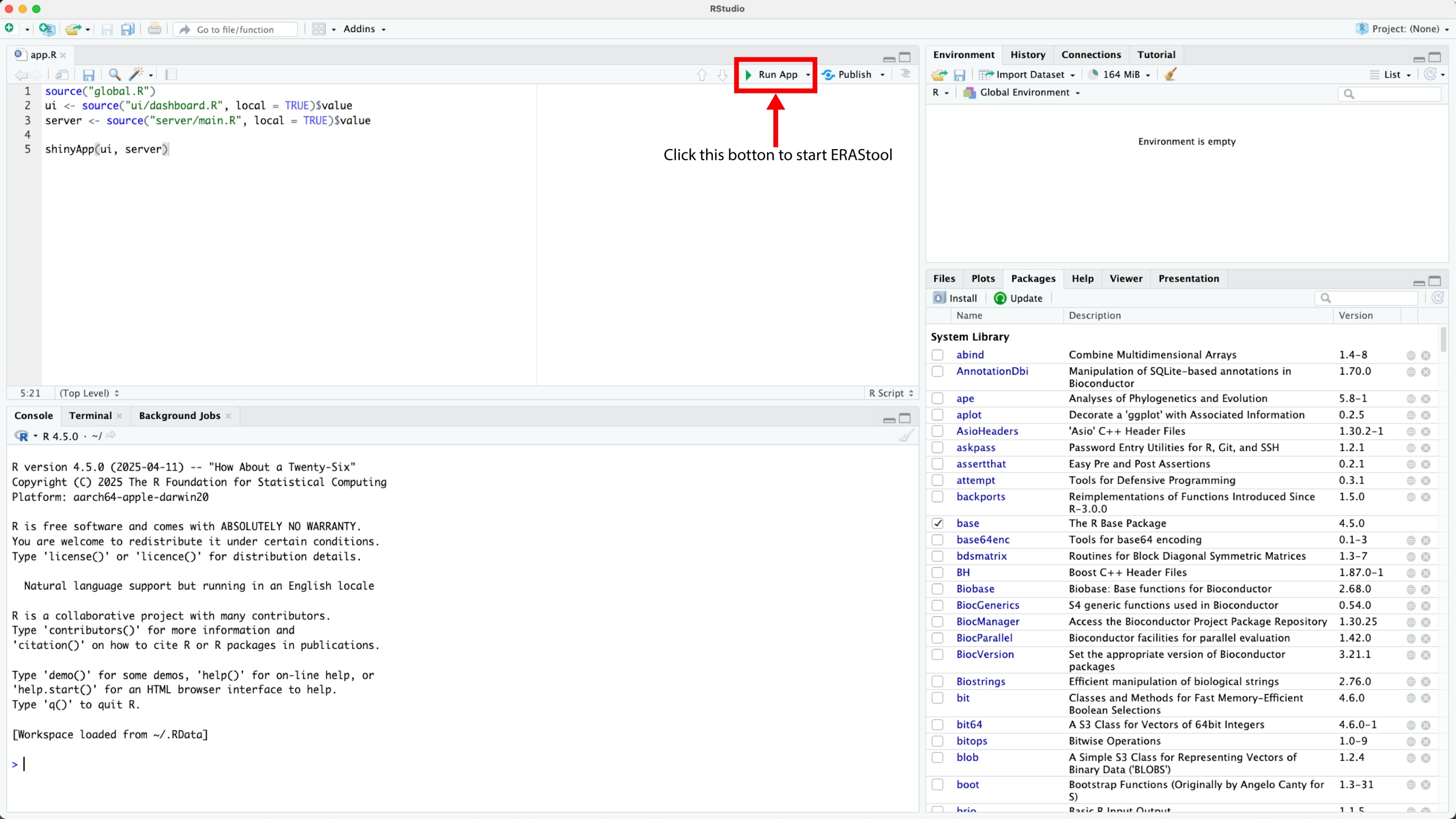

##### Assessing the sample metadata file and the count matrix file to ensure they meet the DESeq2 requirements.

Metadata

File format: .csv or .xlsx

Ensure:

1) Column names must not contain spaces or special characters
2) Row names (sample names) must match exactly with column names in the raw count data
3) Metadata can be directly retrieved from the NIH NICBI GEO depository

Raw Counts

File format: Raw count matrix (.csv, .txt, .xlsx, .gz) or multiple *STAR* .out.tab files

Ensure:

1) Raw count matrix data (not normalized count matrix)
2) Genes are in rows, samples in columns
3) Column names of raw counts = Row names of metadata

#### B. Analytical workflow

1. Adding the student information

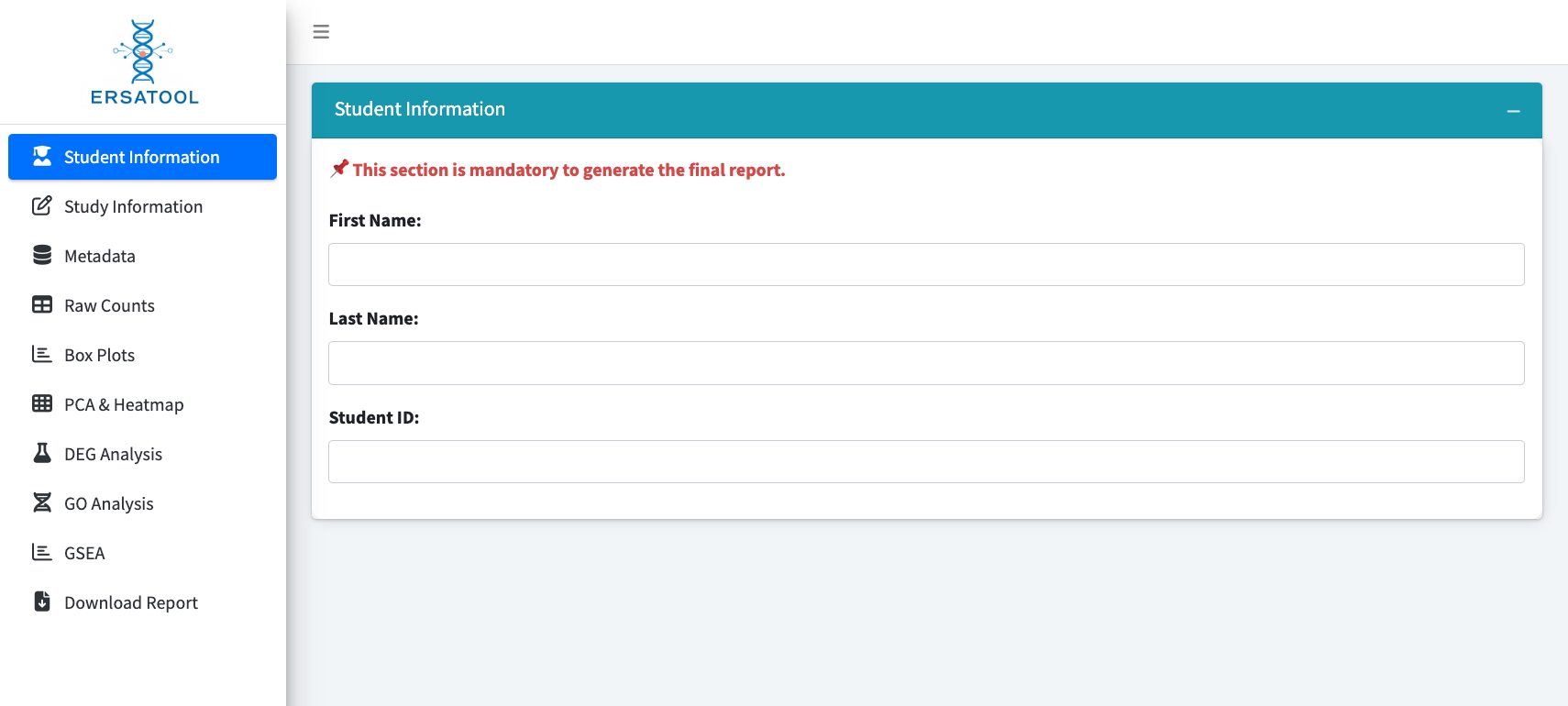

This is the first step of the ERSA tool; these fields are necessary for generating an HTML report file. The HTML report file will be named LastName_date.html.

2. Adding the study information

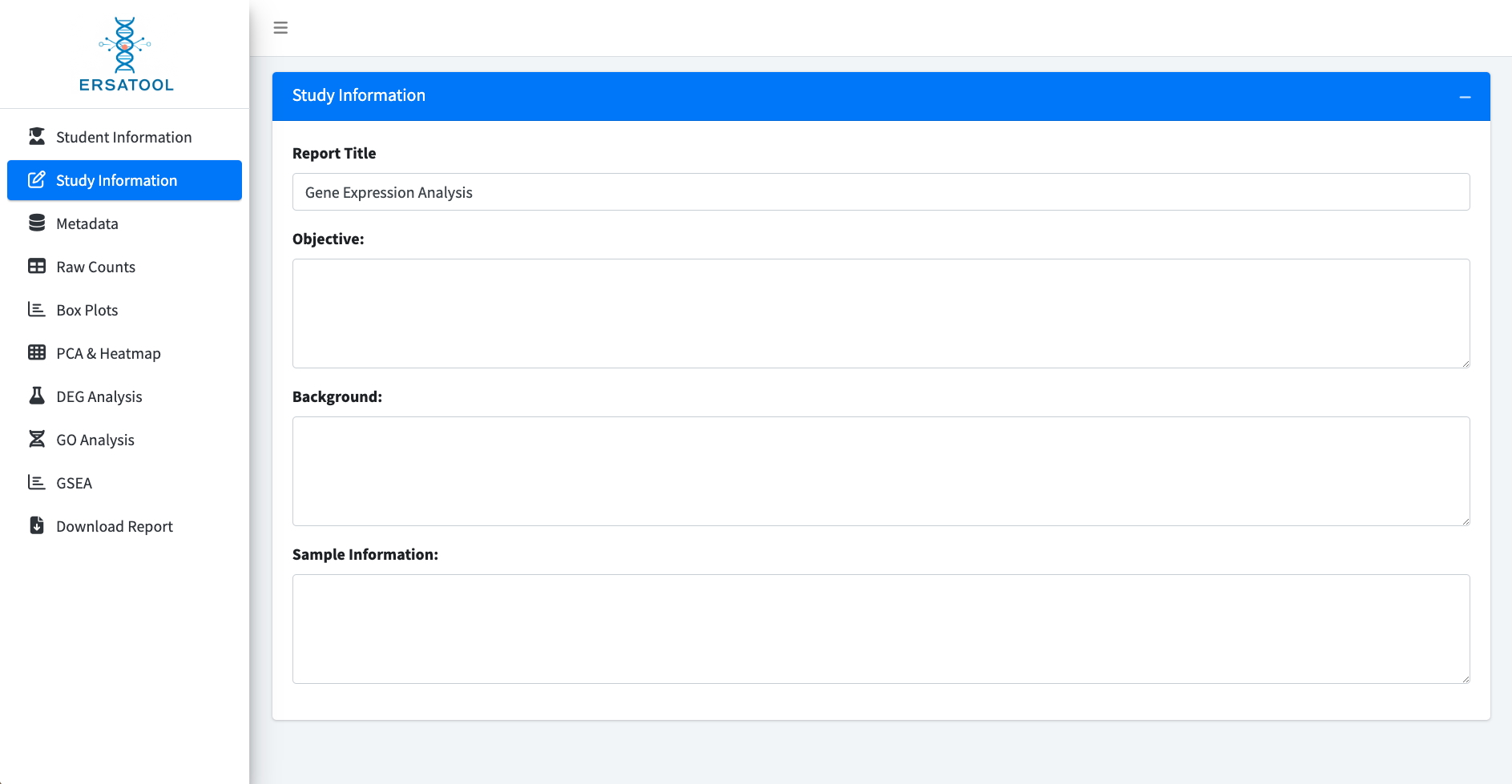

This section has four entry boxes to fill.

1) Report Tile
2) Objective
3) Background
4) Sample information

Although these boxes may remain empty for generating an HTML report file, students are encouraged to fill out all sections to deepen their understanding of the study and clarify the study objective.

3. Loading sample metadata

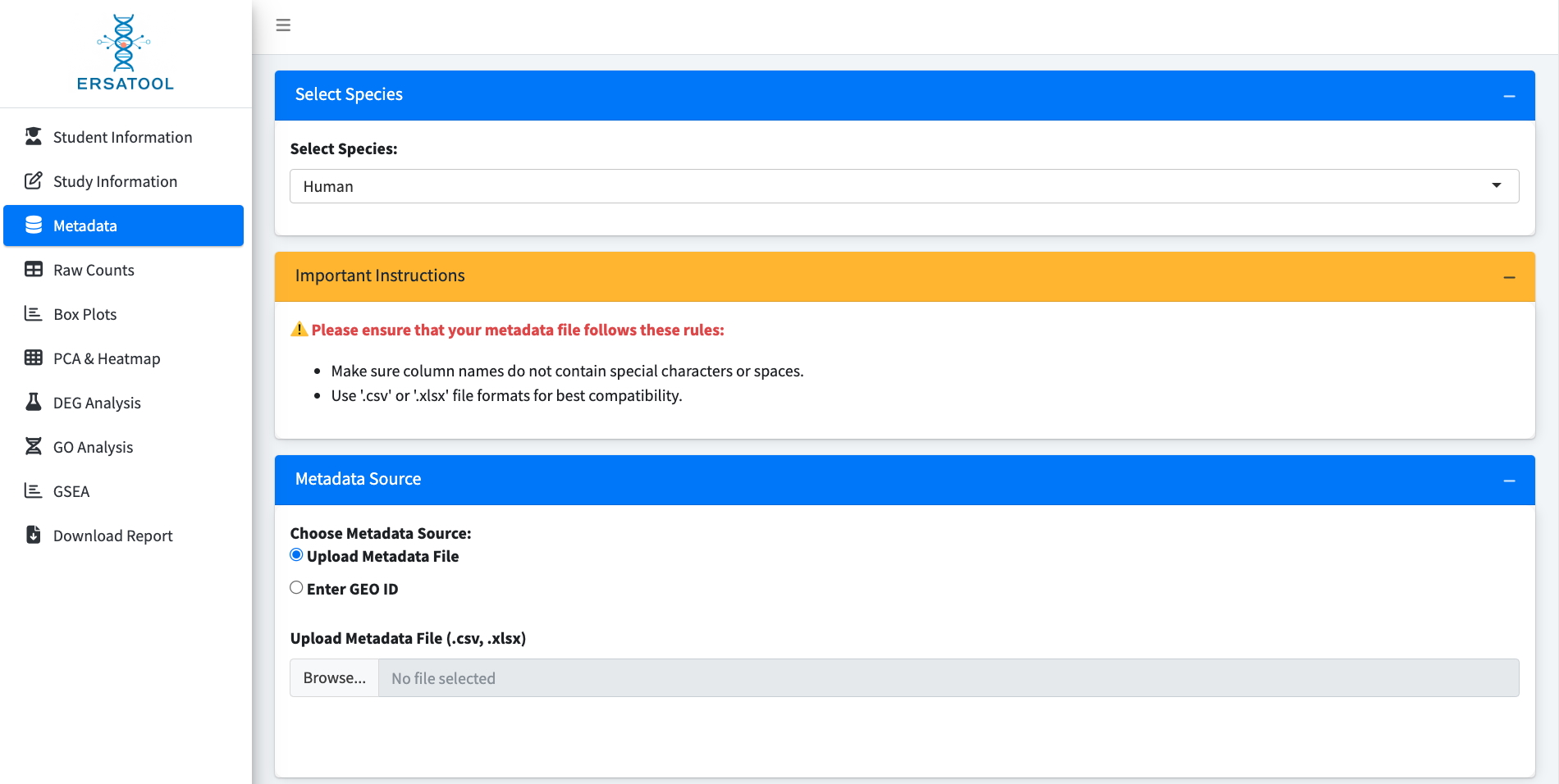

1) Select Species

Currently, the ERSAtool supports analysis for humans and mice.

2) Important Instructions

A few tips to generate a metadata file by the user.

3) Metadata Source

The ERSA tool accepts metadata in .csv and .xlsx file formats and has a function to retrieve sample metadata from the NIH NCBI GEO server. If users obtain sample metadata from GEO, they should ensure that the order of the samples and the names in the metadata and count matrix are consistent.

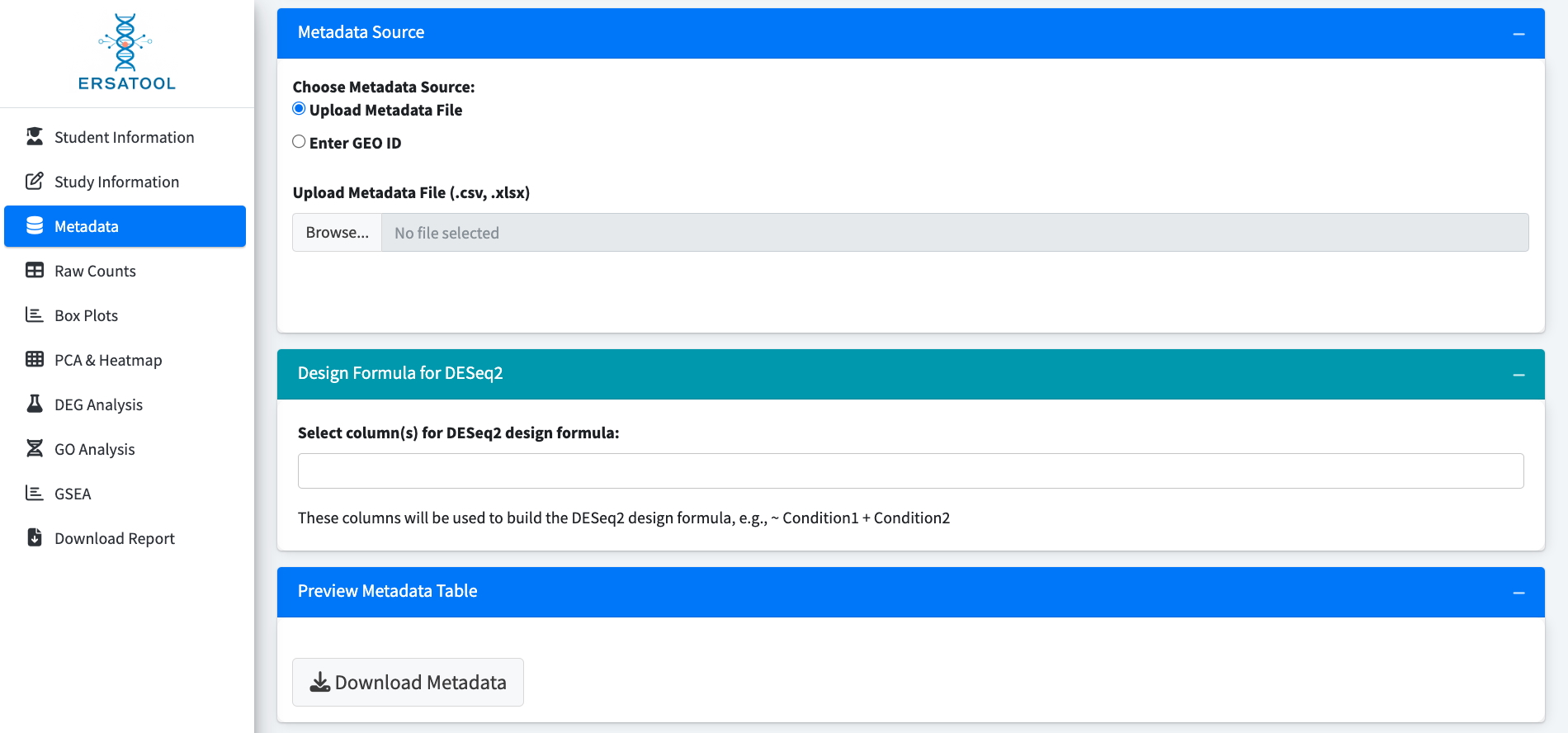

4) Design Formula for DESeq2

Select the column(s) to use for DESeq2 design.

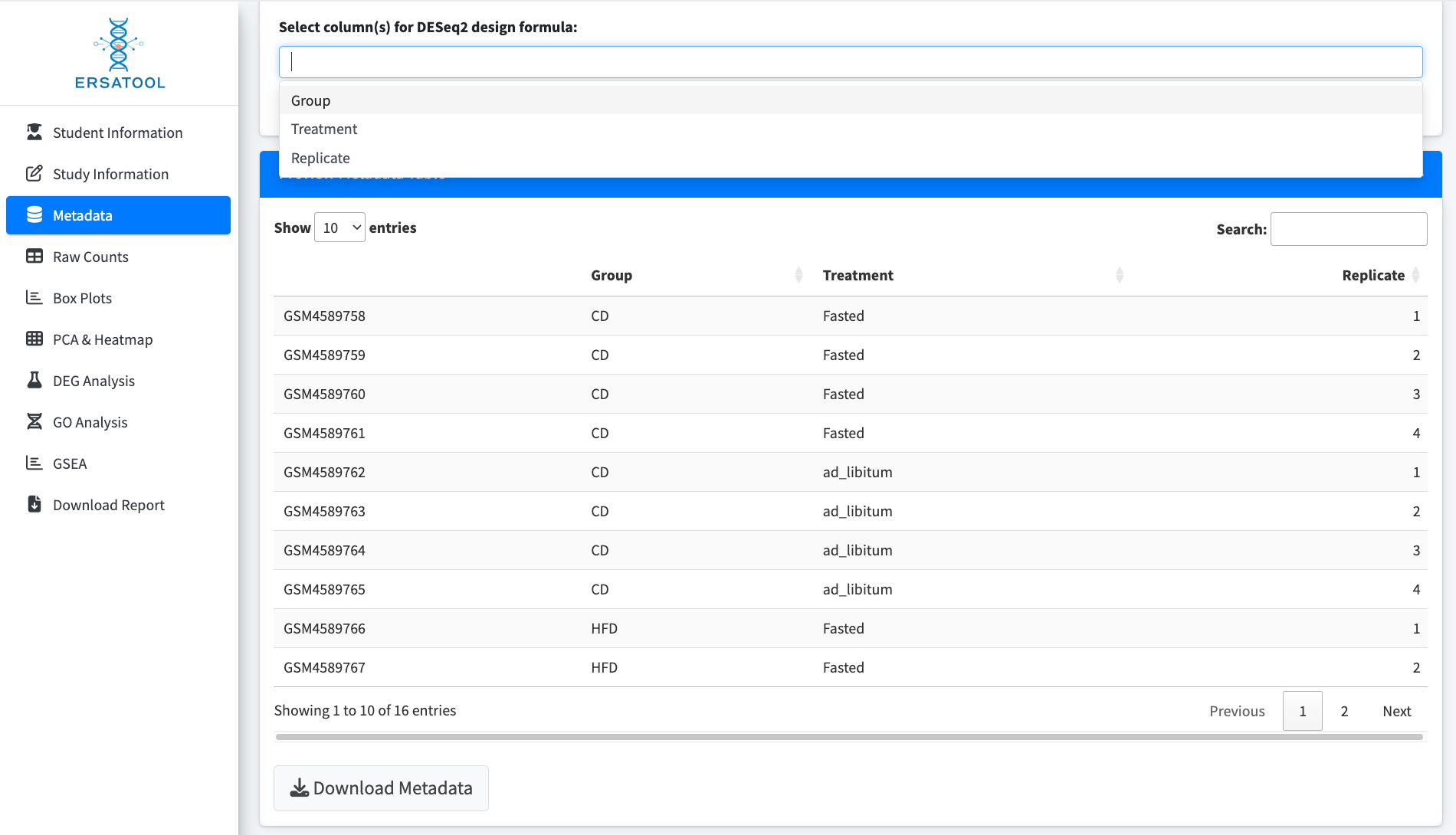

5) Preview Metadata Table

This function allows users to save the edited metadata table for future use or to further modify the metadata (e.g., modifying orders or sample names of the metadata directly retrieved from the GEO) using Excel. The edited metadata file can be re-uploaded as the final metadata file.

6) Modifying the metadata file

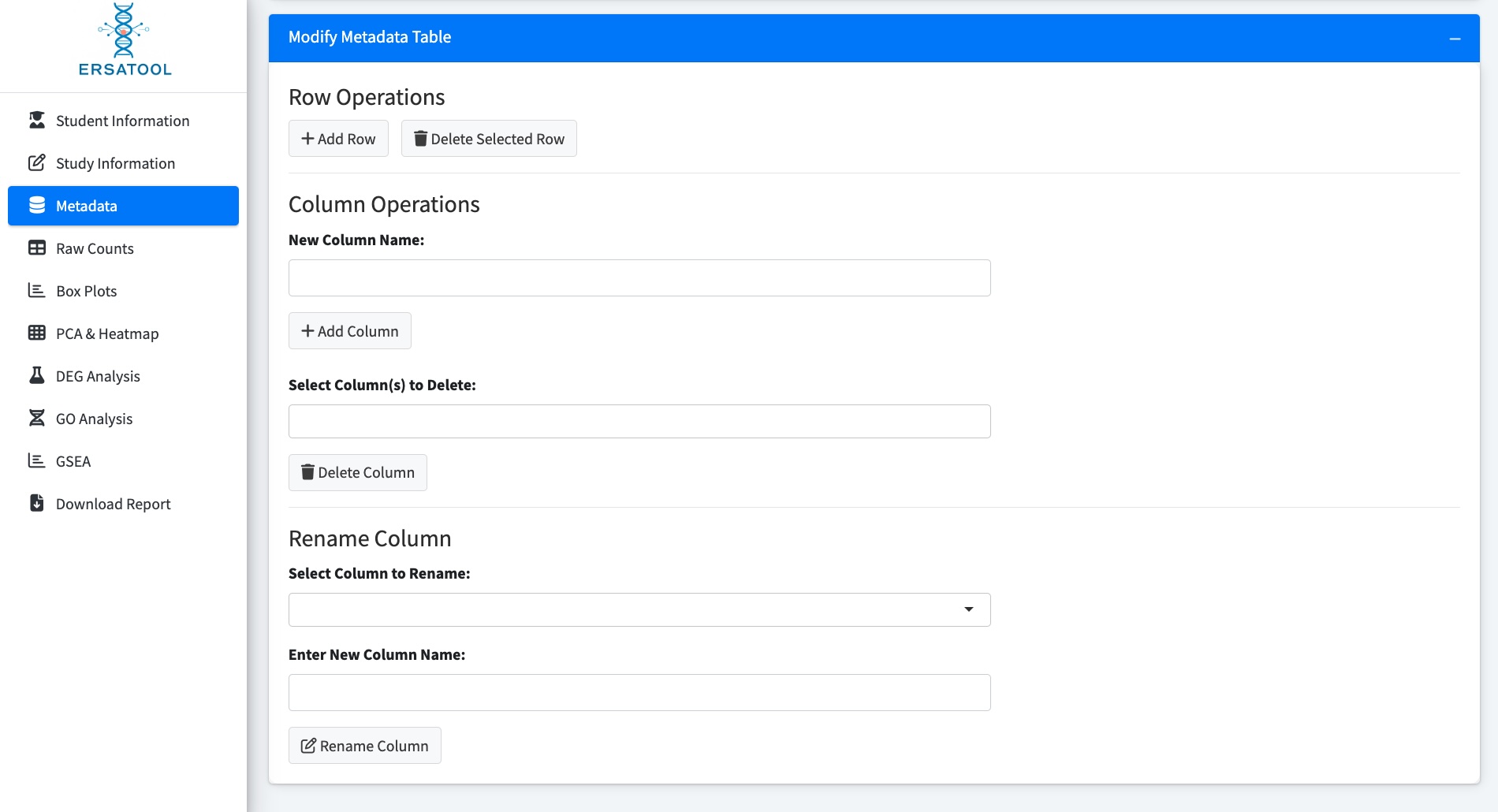

The metadata (rows and columns) can be directly modified on the ERSAtool using the *Modify Metadata Table function*.

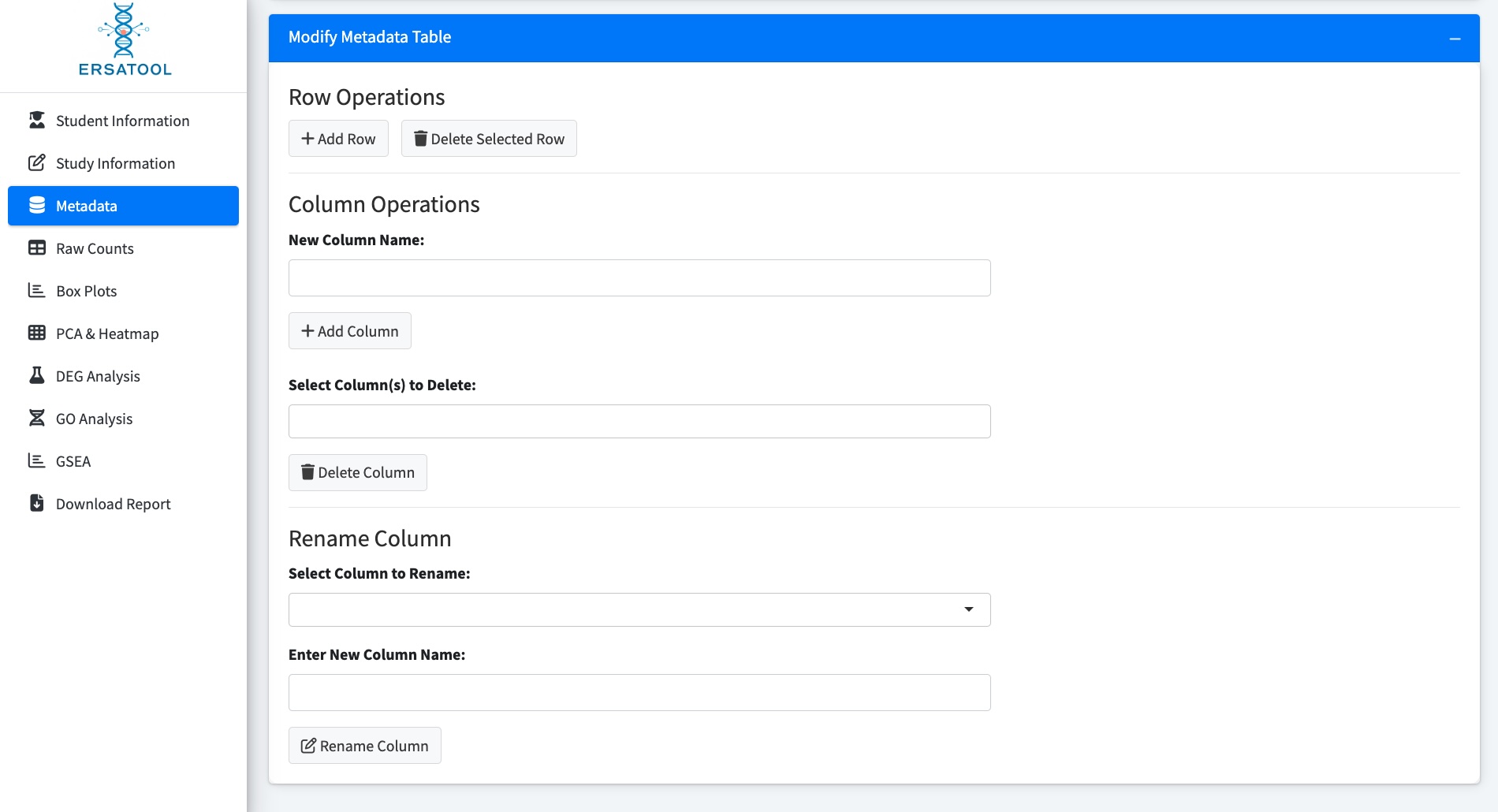

4. Loading raw count matrix

The raw count file can be a raw counts matrix (row names are the gene names or Ensemble IDs, and column names are the sample names) or STAR aligner’s count files (.out.tab).

1) Uploading a raw counts matrix

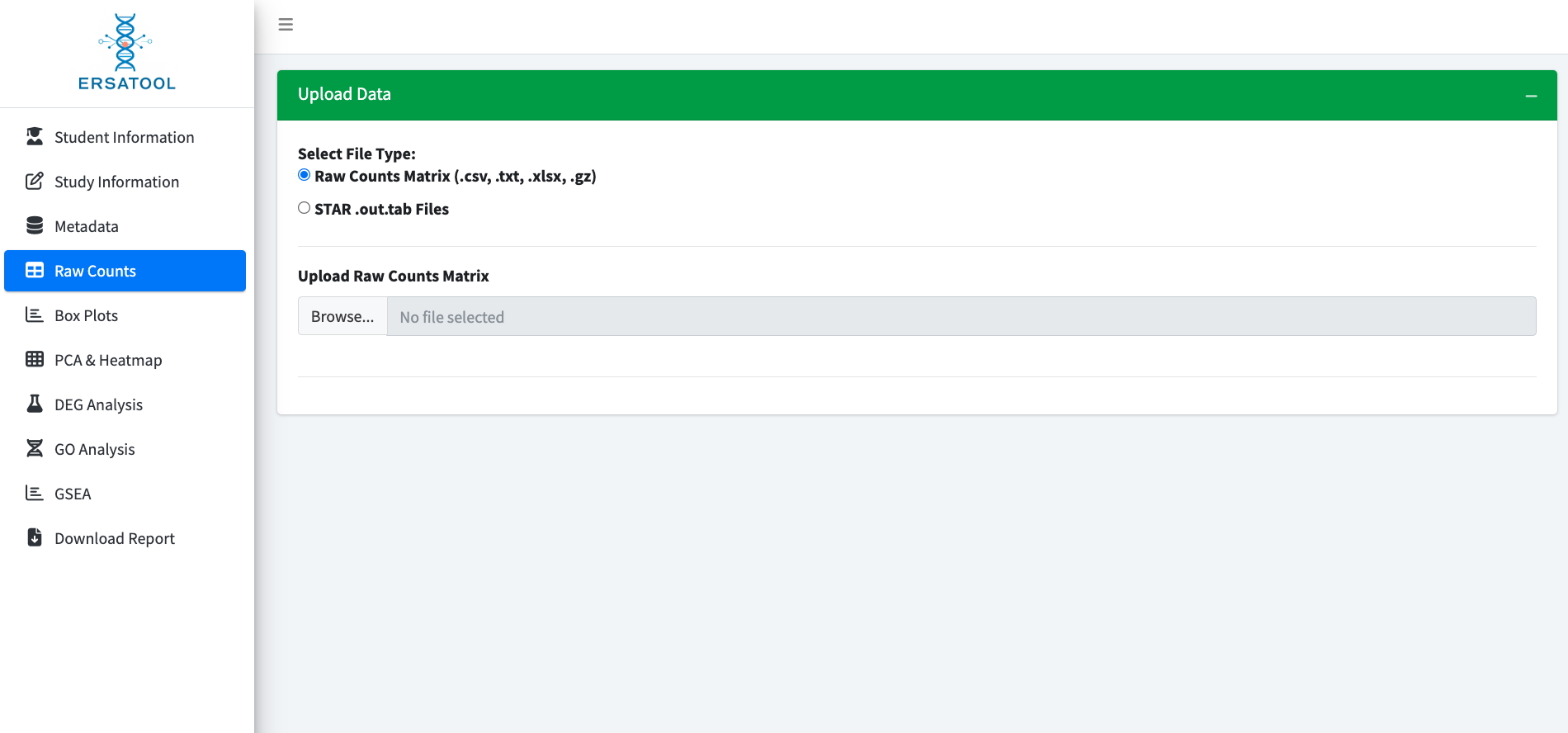

2) Uploading the STAR aligner’s output files
a. Select the column to be used
b. Hit Browse and use the 'Shift’ or 'Command' key to select all .out. tab files to upload.

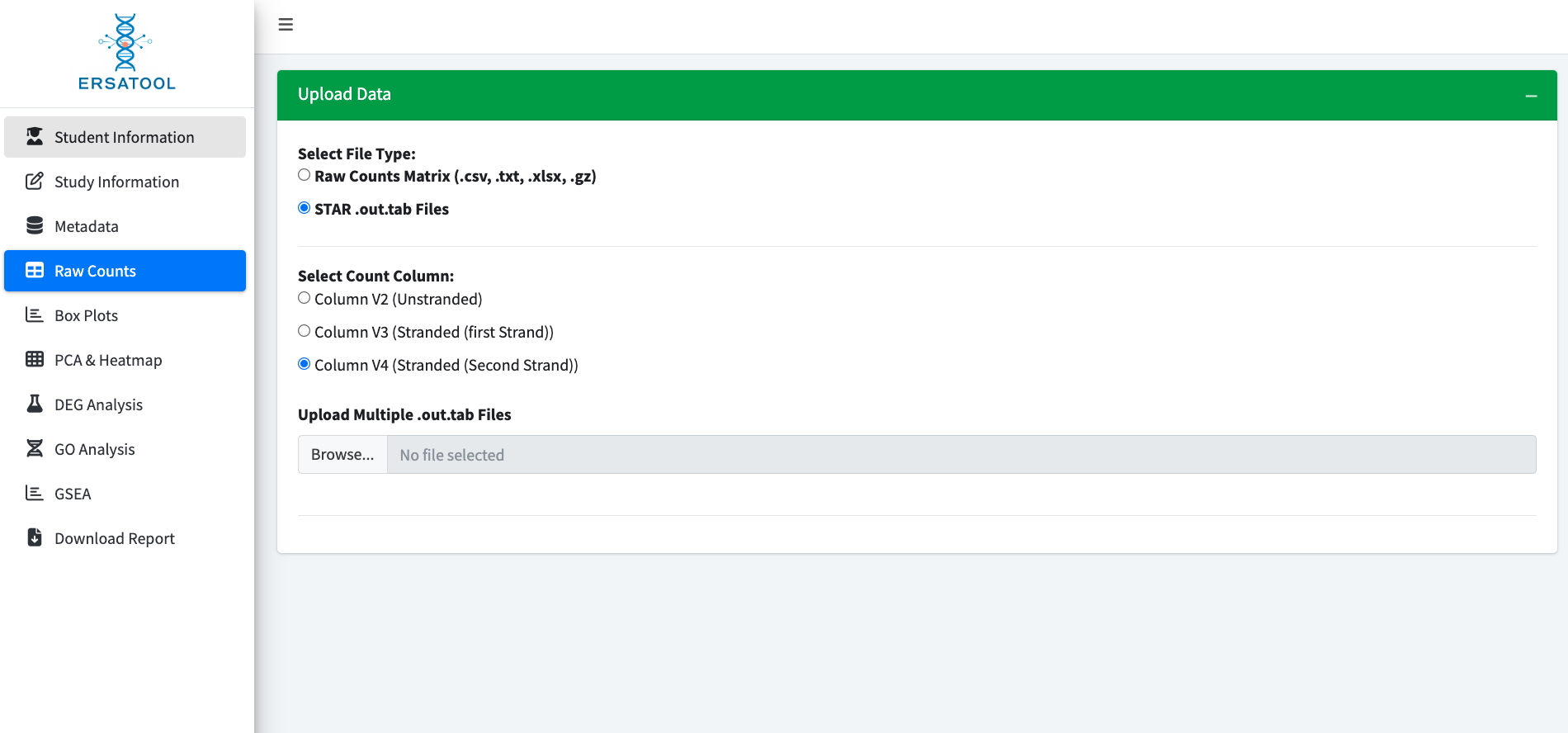

DESeq2 uses raw data for the analysis. If the following error message appears when uploading the count files and the app halts the analysis, it indicates that a normalized count matrix was uploaded.

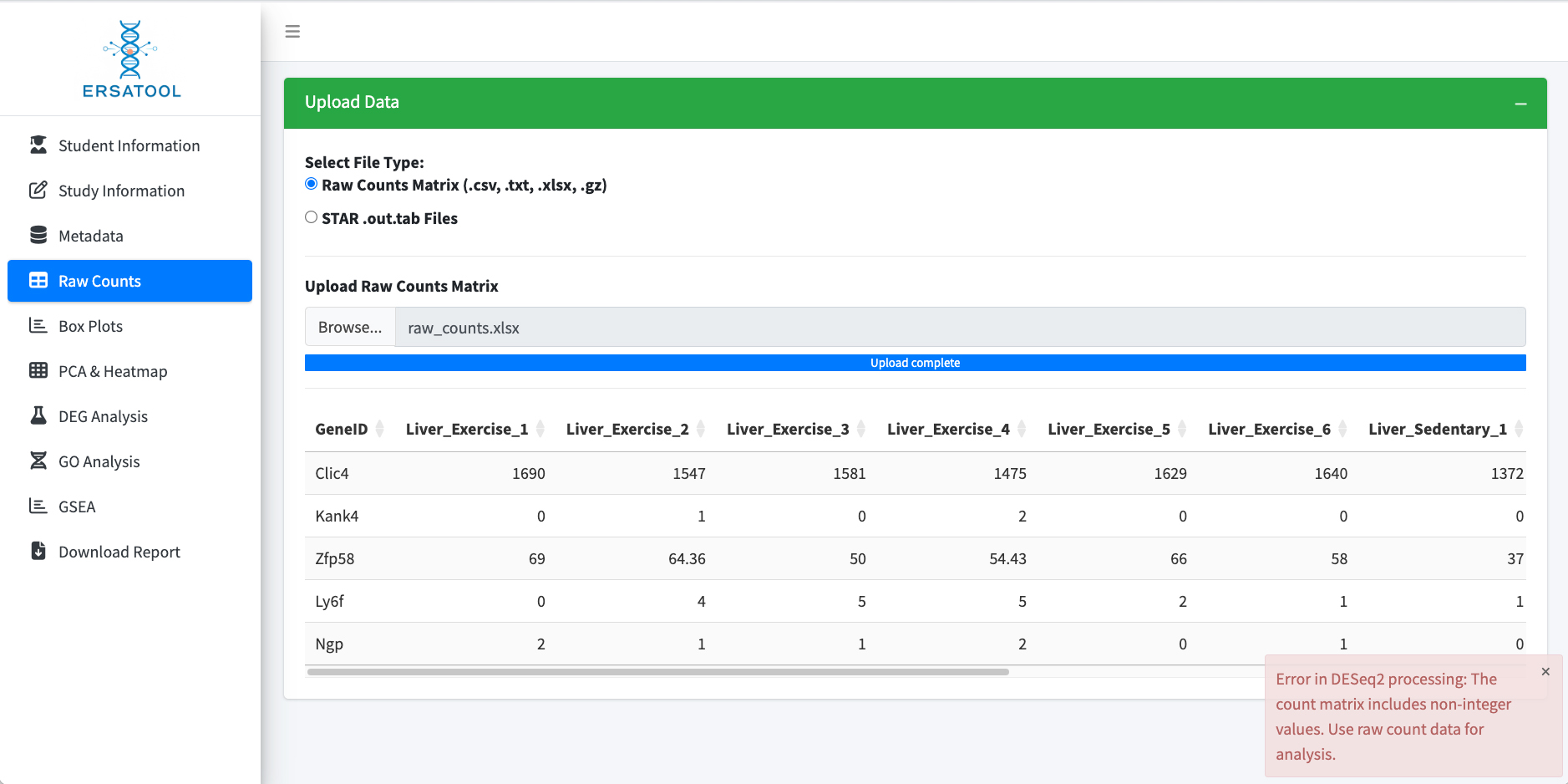

5. Assessing sequencing depth

Here, we displayed the outputs of two datasets, one of which has even count distributions among the samples and the other has uneven distributions. While raw count uneven distribution may not significantly impact the analysis, users must pay attention to these variations when analyzing the datasets.

a) A dataset demonstrating even distributions across the samples

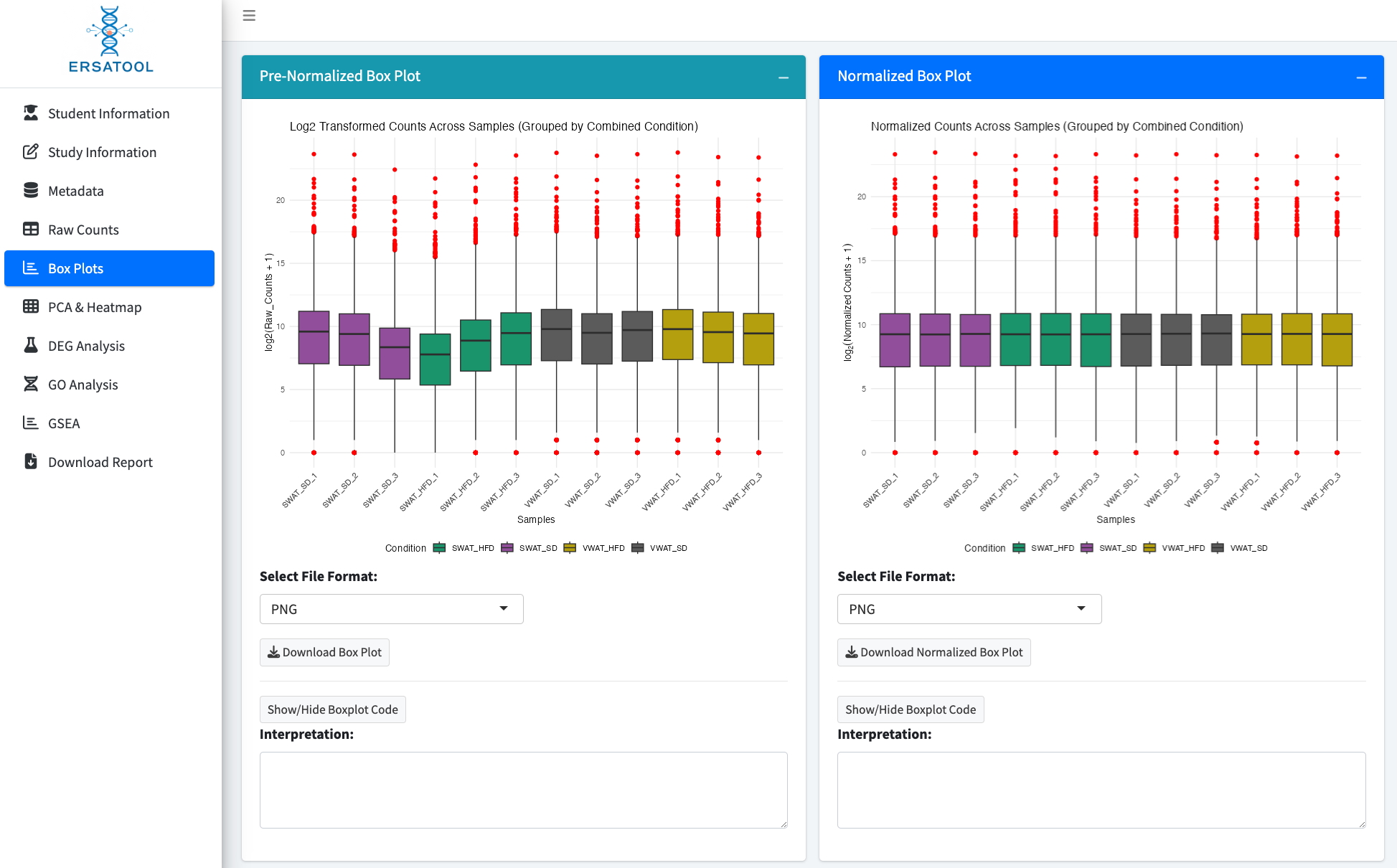

b) A dataset demonstrating uneven distributions across the samples

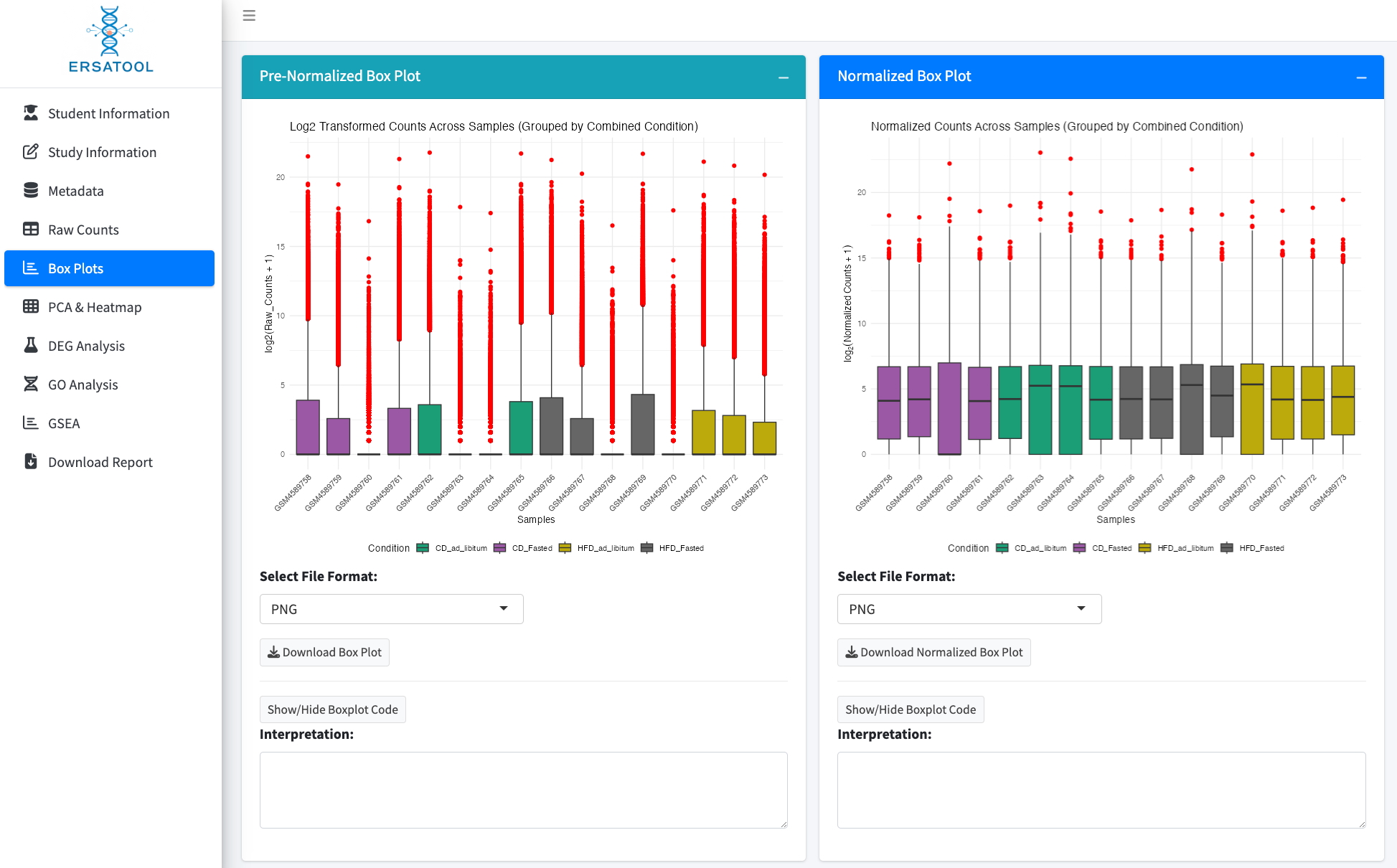

In this dataset, five samples have a very low raw count distribution compared to the other samples. Although the distributions seem improved after normalization, the PCA analysis revealed that PC1, which explained 72.1% of the variance in the difference, separates the samples with higher count distributions from those with lower count distributions. The same was observed on the heatmap using variance-stabilizing transformed values. These results indicate that skewed read distribution contributes to the gene expression profile.

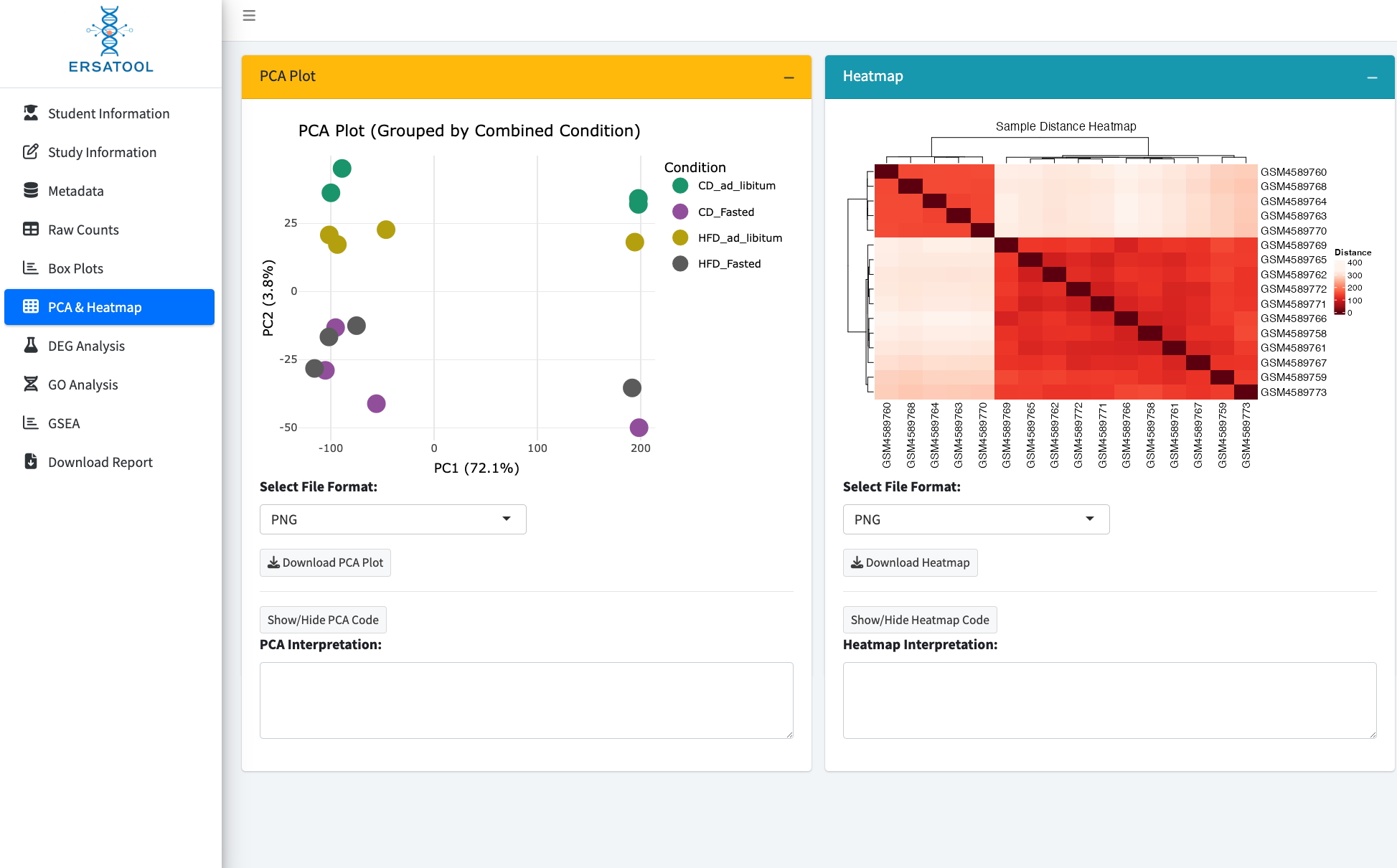

6. Assessing gene expression profile similarity

The ERSAtool generates two exploratory analyses and visualizations to explore sample relationships.

a) Principal component analysis (left).
b) **Heatmap showing distances between samples using variance-stabilizing transformed values (right).**

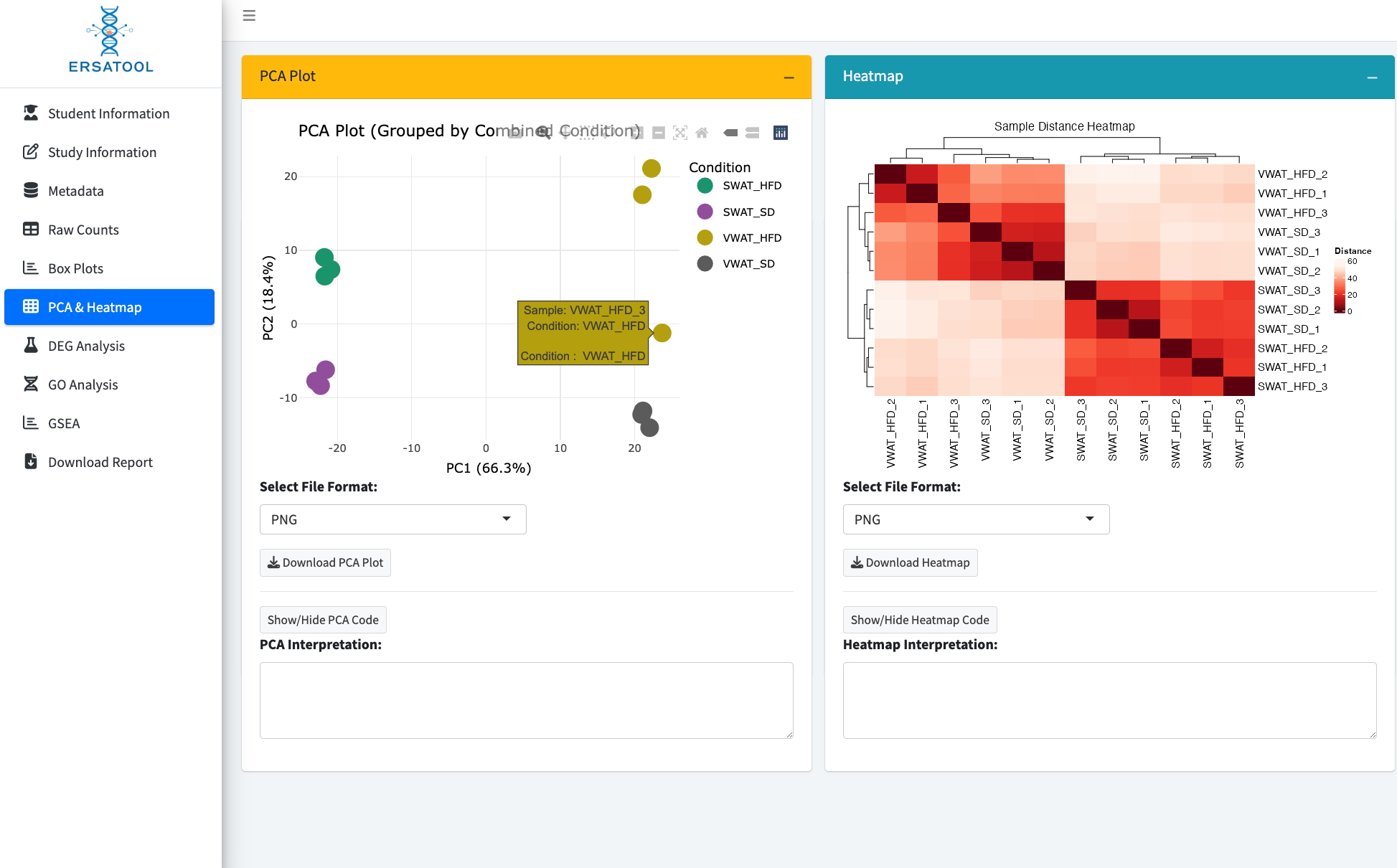

**In this dataset, one sample is identified as an outlier of the group (VWAT_HFD). The user can identify the sample on the PCA plot by hovering the cursor over the marker point.**

7. Identifying and visualizing differentially expressed genes
a) Set-up

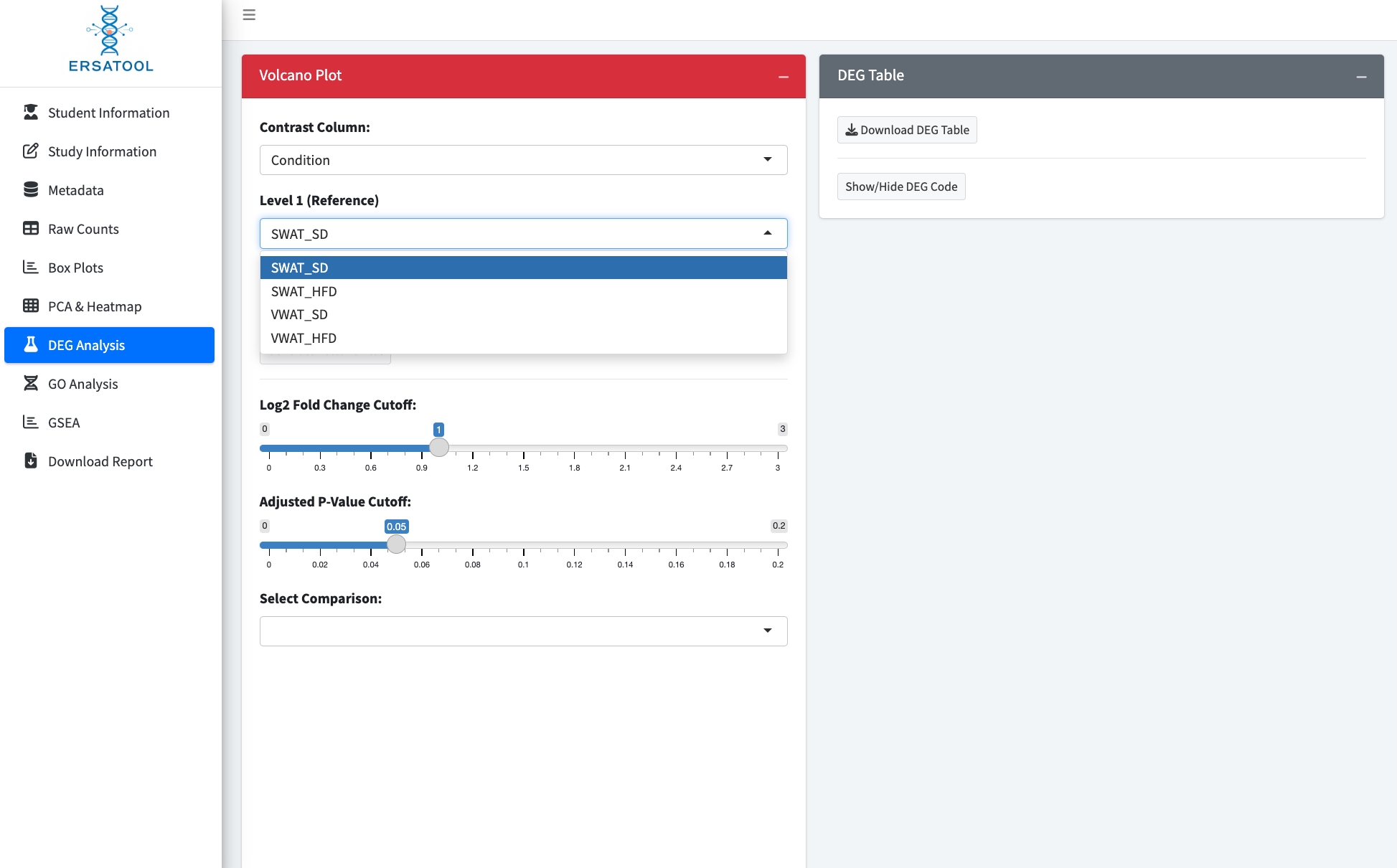

To perform a DEG analysis, select the contrast column and choose comparison pairs from the dropdown menu, then set the thresholds for log2 fold change cutoff and adjusted p-value cutoff. ERSAtool uses xxx as the adjustment method. While the function allows users to set the thresholds, we encourage them to use the default settings (log2 fold change > 1, indicating at least a two-fold difference, and adjusted p-value < 0.05).

b) Generate a volcano plot and a DEG list.

After setting the comparison pairs and threshold, click the “Generate Volcano Plot” button to generate a volcano plot and a DEG list.

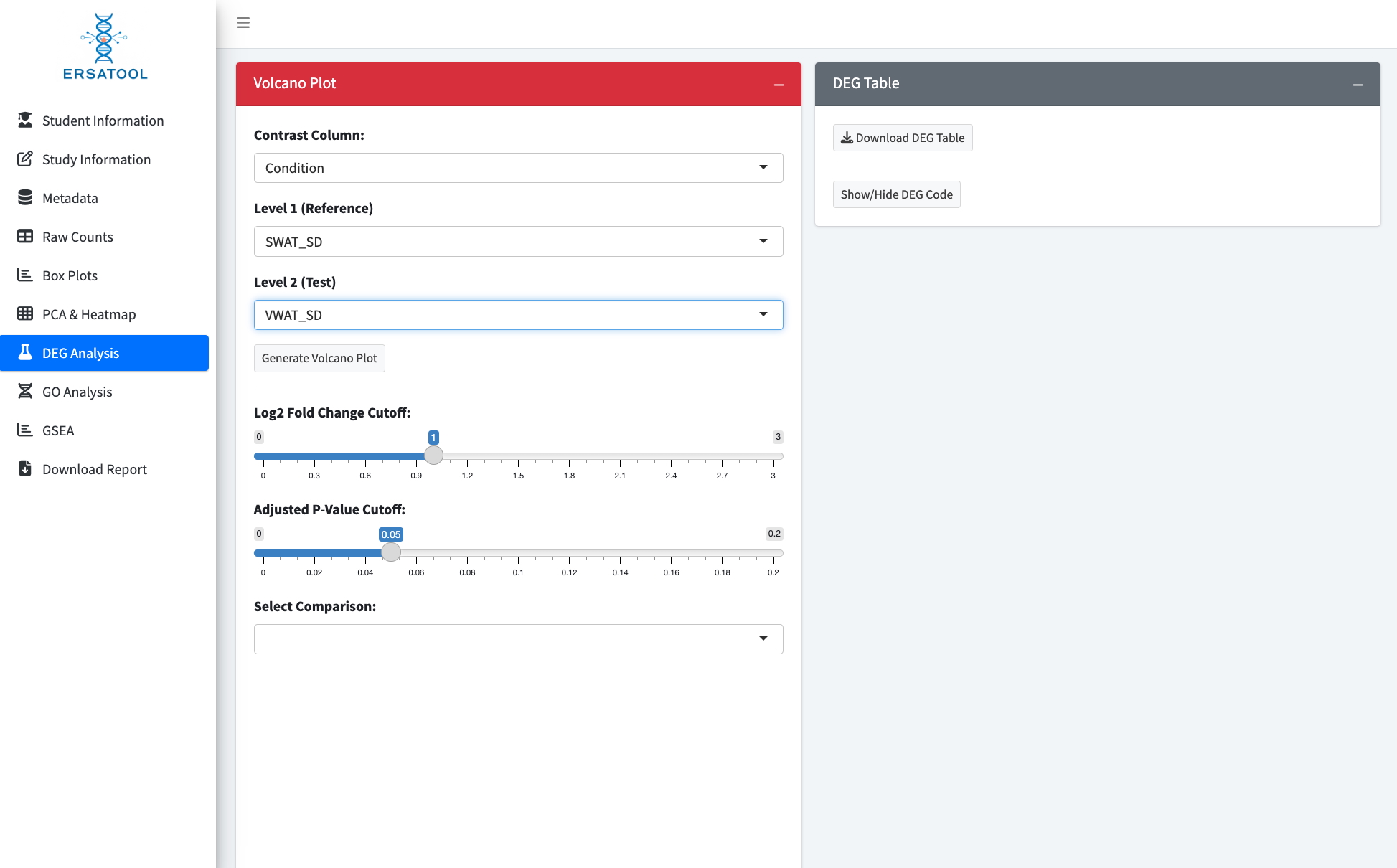

The volcano plot displays the counts of upregulated and downregulated genes, along with the total number of genes tested (a). The top 10 up- and down-regulated genes are labeled on the volcano plot. The search window above the DEG table allows users to check if a specific gene, such as biological marker genes, was identified as a DEG (b).

Note: Whenever users modify the thresholds for the analysis, hit the “Generate Volcano Plot” button again to generate the new results with the updated threshold values.

b. DEG table

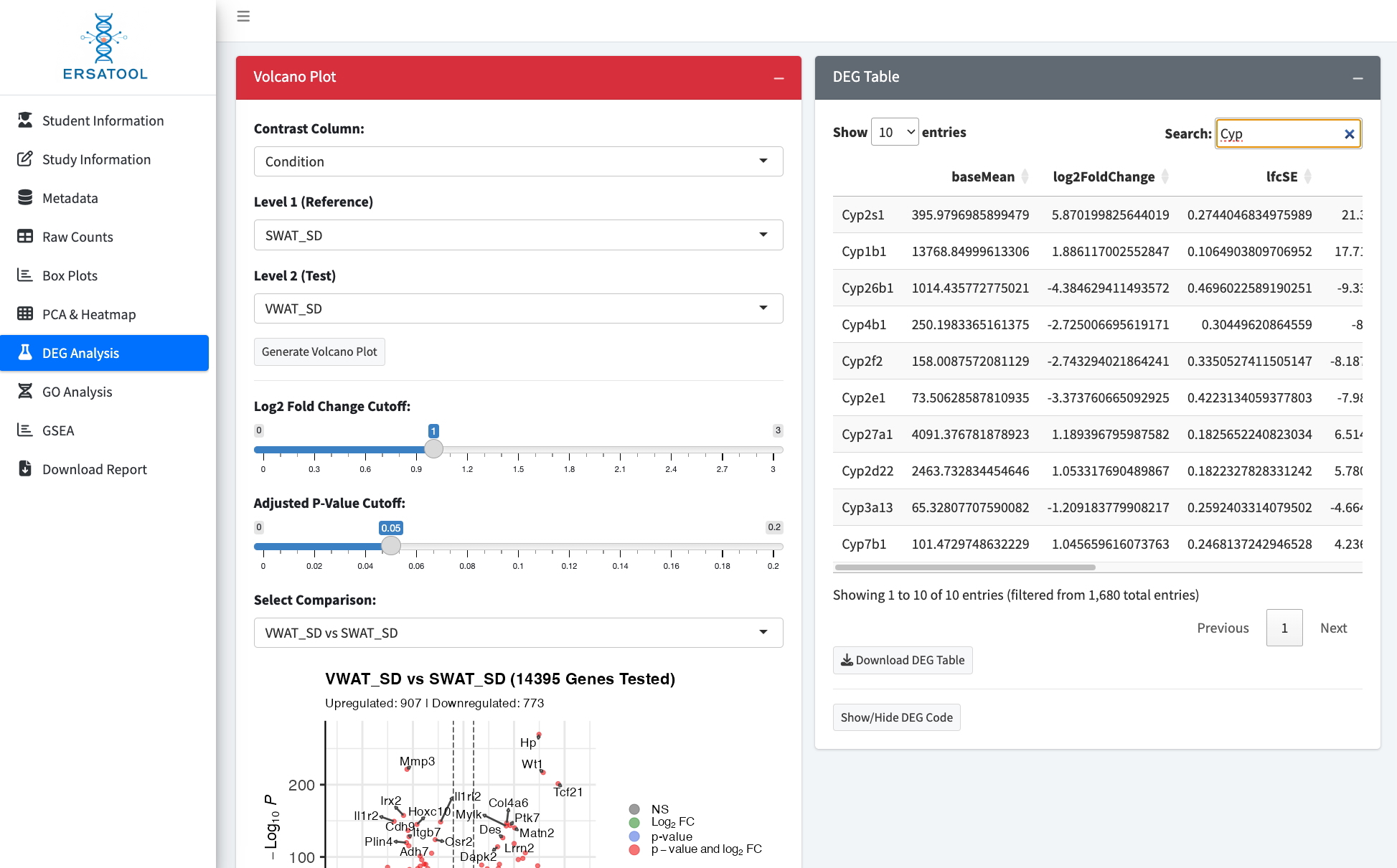

a. Volcano plot

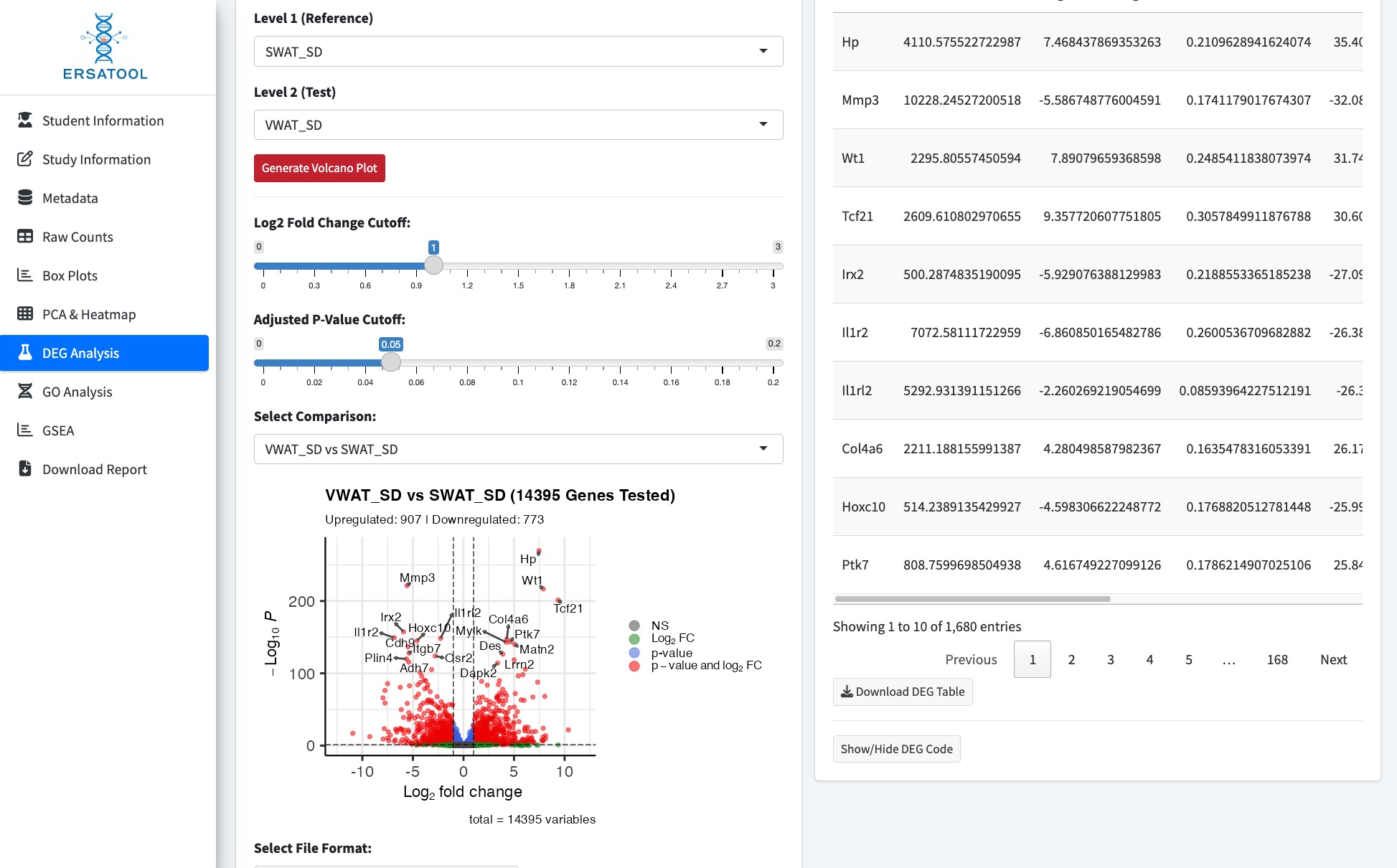

8. Pathway enrichment analysis

ERSAtool provides two types of pathway enrichment analysis: GO enrichment analysis for DEGs (a) and gene set enrichment analysis (GSEA, b). To perform the analysis, hit the “Perform GO Analysis” and “Run GSEA” buttons. ERSAtool performs the analysis using the species user selected above. GO enrichment analysis utilizes the list of DEGs identified in the previous section. Therefore, whenever users adjust the thresholds for the analysis, they should conduct GO enrichment analysis on the newly generated DEGs.

a. GO Analysis

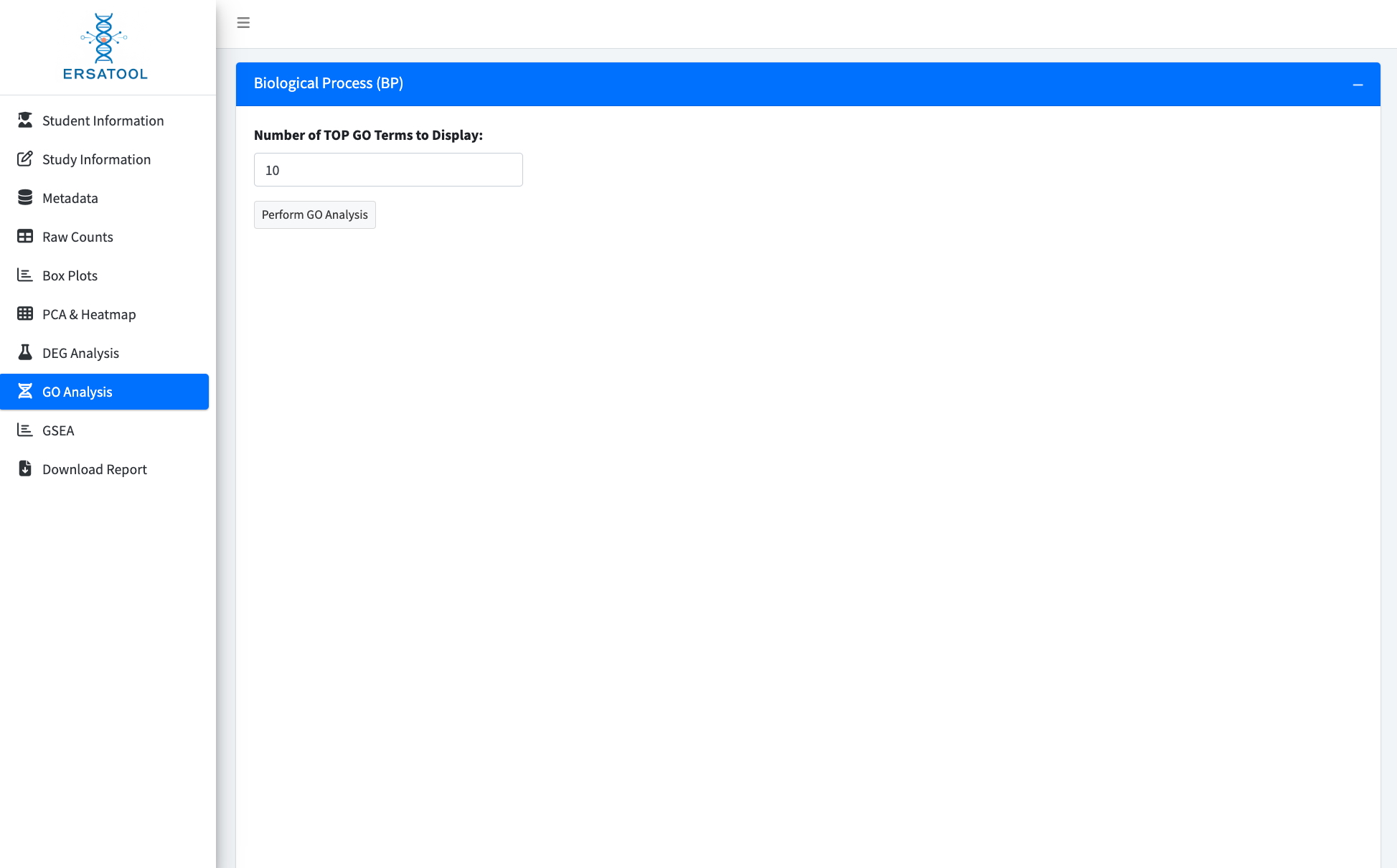

b. GSEA Analysis

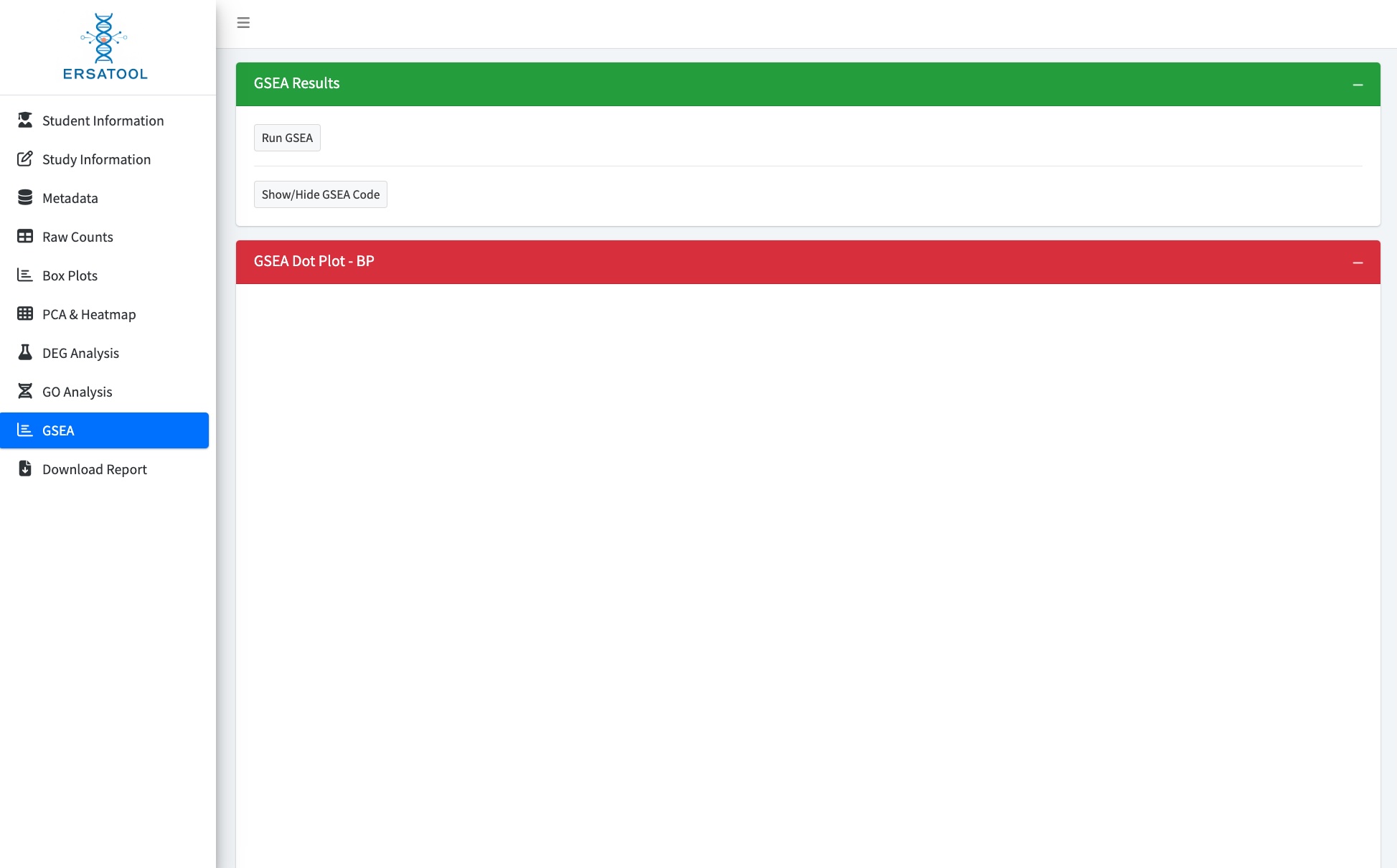

9. Adding a conclusion and exporting analysis results with interpretations as an HTML file

At this step, users add the conclusion or summary to their findings. After that, hit “Generate Report.” The app will consolidate all analysis steps and interpretations added to each step and create a single HTML file for reporting their work.

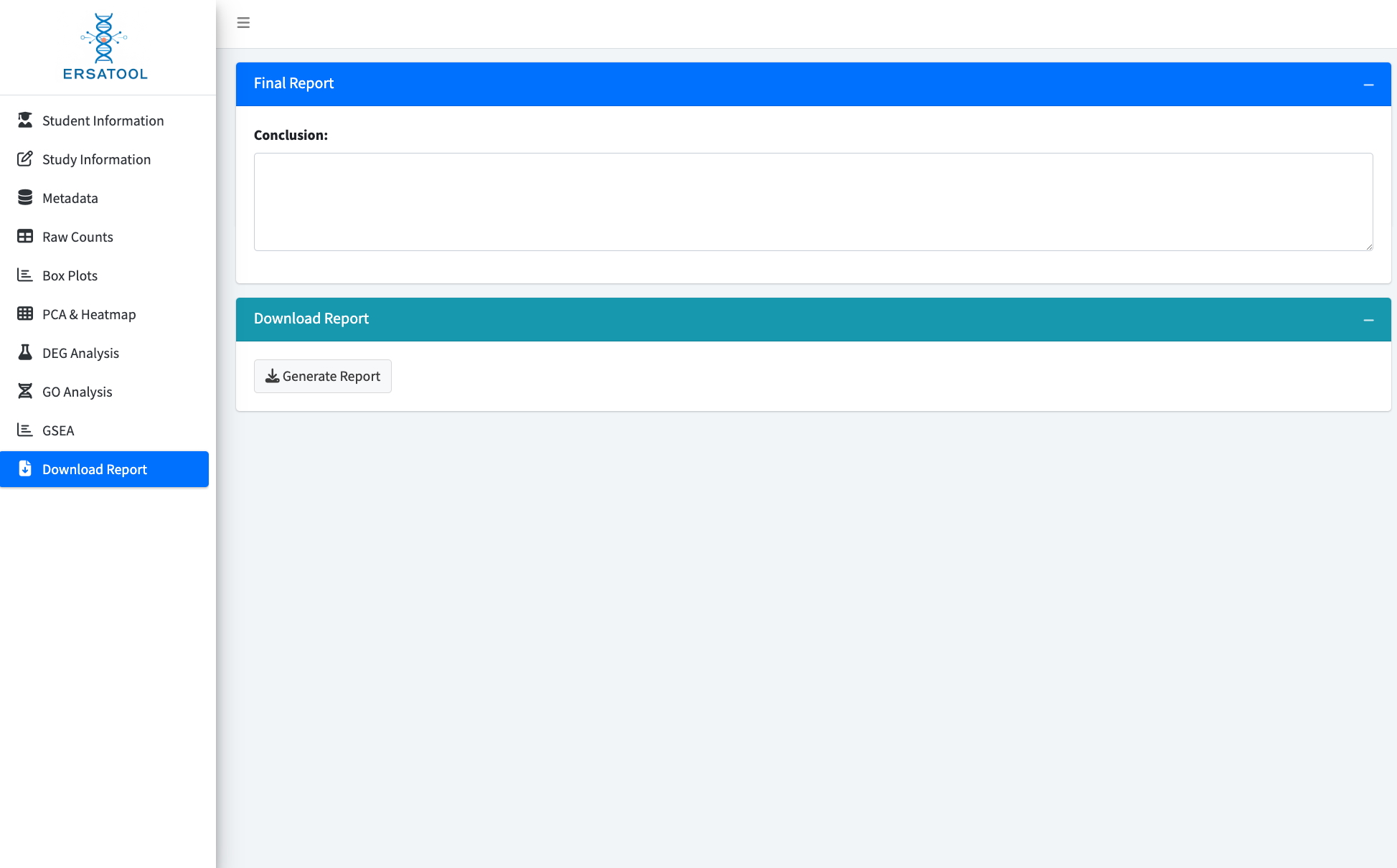

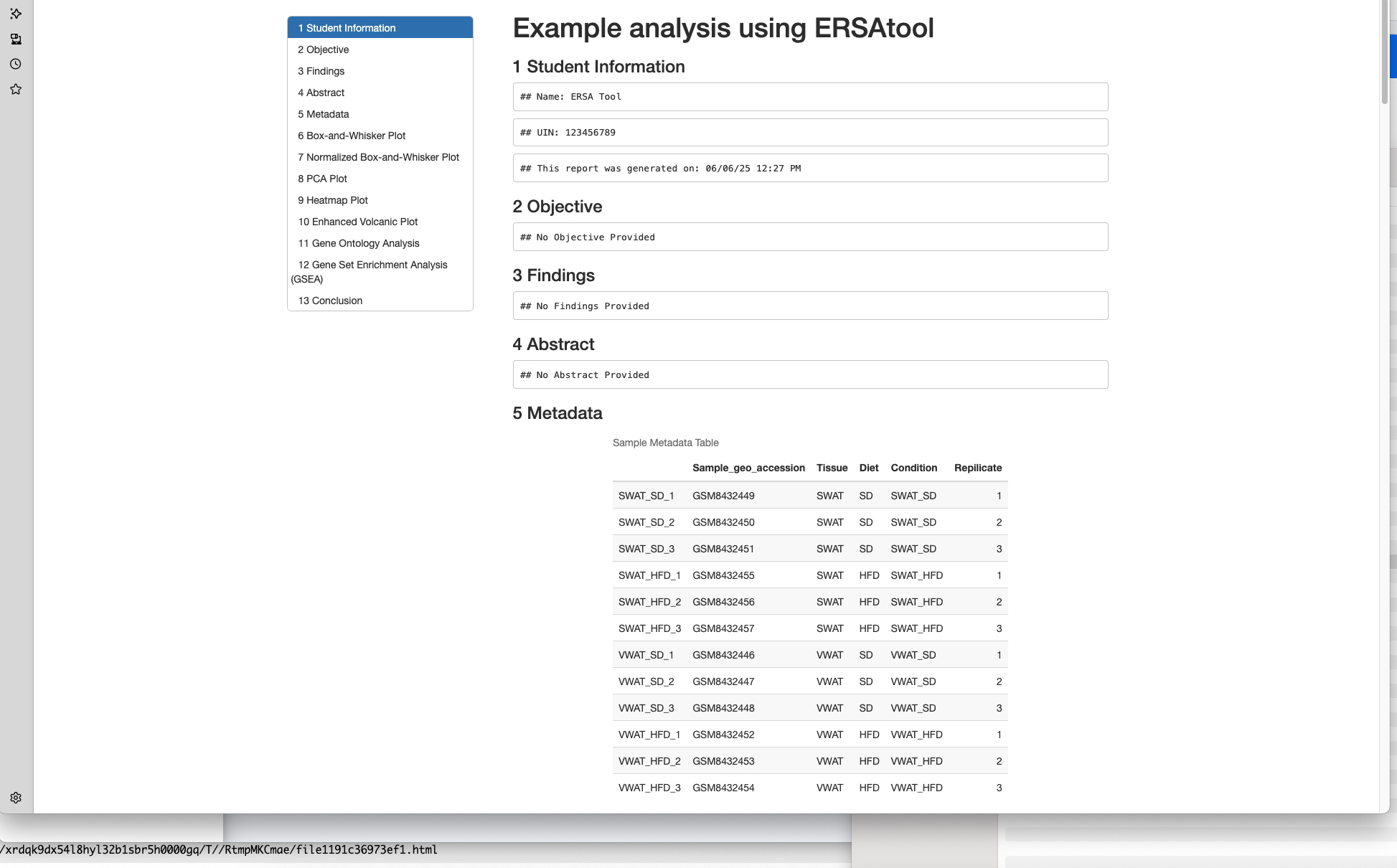

The HTML report file includes a timestamp function to indicate when this report was generated.

10. Useful functions for further learning and reports/publications.

The ERSAtool offers some useful functions for advanced users in addition to generating HTML report files.

1) Showing code used at each step.

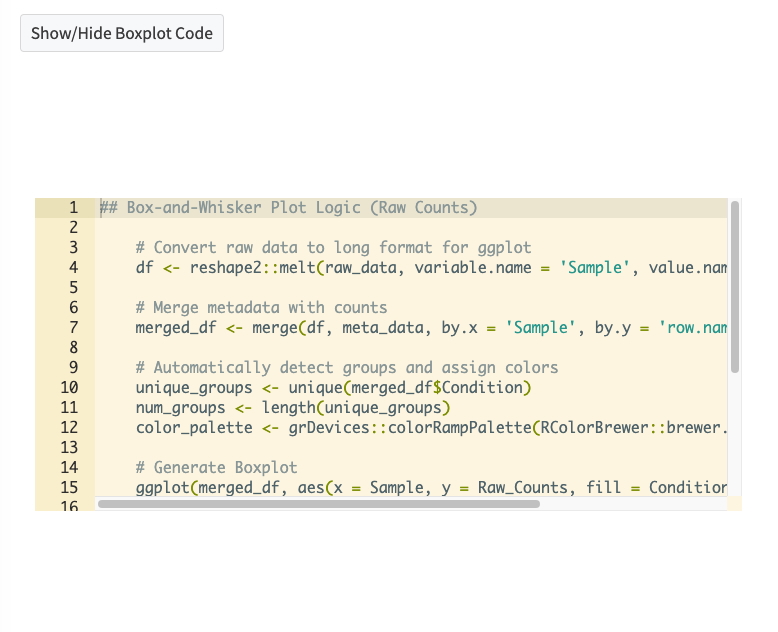

Each section has “Show/hide *** Code” button(s). Simply click them to see the code used in the step.

2) Exporting the plot as a separate image file

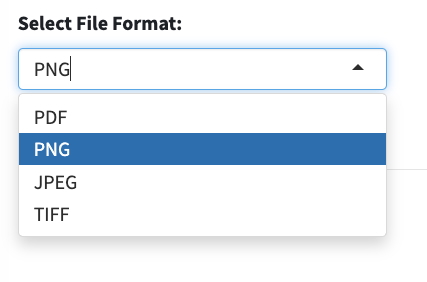

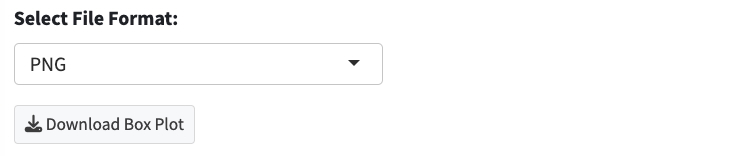

Generated plots can be exported as a single image file (PNG, PDF, JPEG, or TIFF format).
