## Supplementary material for "ERSAtool: A User-Friendly R/Shiny Comprehensive Transcriptomic Analysis Interface Suitable for Education": Tool_06_13_25.html

Changes in tissue-specific gene expression of adipose precursor cells due to high-fat diet intake


### Changes in tissue-specific gene expression of adipose precursor cells due to high-fat diet intake

### 1 Student Information

```
## Name: Ersa Tool
```

```
## UIN: 123456789
```

```
## This report was generated on: 06/13/25 02:55 PM
```

### 2 Objective

```
## The aim of this case study is reviewing the gene expression alterations between adipocyte precursor cells (APCs) from subcutaneous white adipose tissue (SWAT) and visceral white adipose tissue (VWAT), which were sorted from mice fed a 60% kcal lard high-fat diet (HFD) for 3 days, a period when APC proliferation peaks.
```

### 3 Findings

```
## Adipocyte hyperplasia is a significant mechanism of adipose expansion. The original study found that HFD feeding accelerated proliferation in a tissue-specific manner. Additionally, the authors observed that the tissue-specific alteration occurred only in males. We aim to assess whether a high-fat diet changes the expression of genes associated with cell cycle pathways in male APCs derived from different fat tissues.
```

### 4 Abstract

```
## We have 12 samples in total from four groups (two tissue types and two diet types). All samples are from male mice.
## SWAT-SD (n=3): APC isolated from SWAT from mice fed a standard diet (SD)
## SWAT-HFD (n=3): APC isolated from SWAT from mice fed an HFD
## VWAT-SD (n=3): APC isolated from VWAT from mice fed a standard diet (SD)
## VWAT-HFD (n=3): APC isolated from VWAT from mice fed an HFD
```

### 5 Metadata

Sample Metadata Table

|  | Sample\_geo\_accession | Tissue | Diet | Condition | Repilicate |
| --- | --- | --- | --- | --- | --- |
| SWAT\_SD\_1 | GSM8432449 | SWAT | SD | SWAT\_SD | 1 |
| SWAT\_SD\_2 | GSM8432450 | SWAT | SD | SWAT\_SD | 2 |
| SWAT\_SD\_3 | GSM8432451 | SWAT | SD | SWAT\_SD | 3 |
| SWAT\_HFD\_1 | GSM8432455 | SWAT | HFD | SWAT\_HFD | 1 |
| SWAT\_HFD\_2 | GSM8432456 | SWAT | HFD | SWAT\_HFD | 2 |
| SWAT\_HFD\_3 | GSM8432457 | SWAT | HFD | SWAT\_HFD | 3 |
| VWAT\_SD\_1 | GSM8432446 | VWAT | SD | VWAT\_SD | 1 |
| VWAT\_SD\_2 | GSM8432447 | VWAT | SD | VWAT\_SD | 2 |
| VWAT\_SD\_3 | GSM8432448 | VWAT | SD | VWAT\_SD | 3 |
| VWAT\_HFD\_1 | GSM8432452 | VWAT | HFD | VWAT\_HFD | 1 |
| VWAT\_HFD\_2 | GSM8432453 | VWAT | HFD | VWAT\_HFD | 2 |
| VWAT\_HFD\_3 | GSM8432454 | VWAT | HFD | VWAT\_HFD | 3 |

### 6 Box-and-Whisker Plot

#### 6.1 Box-and-Whisker Interpretation

```
## The box plot indicates the distributions of raw counts among the samples. The upper edge of the box is the 75th percentile, the lower edge represents the 25th percentile, and the horizontal line in the box indicates the 50th percentile. Two samples, SWAT_SD_3 and SWAT_HFD_1, showed lower raw count distributions.
```

### 7 Normalized Box-and-Whisker Plot

#### 7.1 Normalized Box-and-Whisker Interpretation

```
## While I observed that some samples have lower raw count distributions, all showed similar count distributions after normalization using library sizes.
```

### 8 PCA Plot

#### 8.1 PCA Interpretation

```
## Principal component analysis (PCA) visualizes sample similarities in reduced dimensions. In this plot, we displayed PC1, which accounts for 66.3% of the variation, on the x-axis and PC2, which accounts for 18.4% of the variation, on the y-axis. Except for one sample, VWAT_HFD_3, I observed significant segregation among groups. Tissue differences (SWAT or VWAT) were associated with PC1, while diet differences (SD or HFD) were linked to PC2.
```

### 9 Heatmap Plot

#### 9.1 Heatmap Interpretation

```
## The heatmap utilized the rlog-transformed values. The hierarchical dendrogram demonstrates the similarity of gene expression profiles between the samples. Similar to our observations in the PCA plot, I noted a distinct branching between SWAT and VWAT, with SD and HFD branching further among the SWAT samples.
```

### 10 Enhanced Volcanic Plot

#### 10.1 Volcanic Plot Interpretation

```
## In this analysis, I examined the differences in 14,395 genes. My comparison was SWAT_HFD (reference) and VWAT_HFD (test). I identified differentially expressed genes using an adjusted p-value of <0.05 and a log2 fold change greater than 0.6 (which is equivalent to at least 1.5 times the difference between the groups). I found that 1374 genes were upregulated and 1291 were downregulated in VWAT_HFD compared to SWAT_HFD.
```

### 11 Gene Ontology Analysis

#### 11.1 Biological Process

##### Biological Process Plot Interpretation

```
## The GO enrichment analysis tested if DEGs are enriched in specific pathways. Based on this analysis, I found that VWAT_HFD increased genes in nuclear divisions and sister chromatid segregation related pathways and decreased expression in immune cell-related pathways.
```

#### 11.2 Molecular Function

##### 11.2.1 Molecular Function Plot Interpretation

```
## The identified enriched MF GO terms are comparable to those of BP GO terms.
```

### 12 Gene Set Enrichment Analysis (GSEA)

#### 12.1 GSEA Dot Plot - Biological Process (BP)

#### 12.2 GSEA BP Plot Interpretation

```
## No GSEA BP Plot Interpretation Provided
```

#### 12.3 GSEA Dot Plot - Molecular Function (MF)

#### 12.4 GSEA MF Plot Interpretation

```
## No GSEA MF Plot Interpretation Provided
```

### 13 Conclusion

```
## This case study examines gene expression changes in adipocyte precursor cells (APCs) from subcutaneous (SWAT) and visceral (VWAT) white adipose tissues of male mice on a 60% kcal lard HFD for 3 days during peak APC proliferation. We analyzed 12 samples across four groups (two tissue types and two diet types), with all samples derived from male mice. Although two samples had slightly lower distributions before normalization, all samples exhibited similar counts afterward. Therefore, all samples possess comparable sequence depth and are suitable for comparison.
## Principal component analysis (PCA) indicated that tissue differences align with principal component 1 (66.3% variation), while diet differences correspond with principal component 2 (18.4% variation). A heatmap utilizing rlog-transformed values illustrated that, similar to the PCA findings, there was distinct branching between SWAT and VWAT, with SD and HFD branching further among SWAT samples. These results suggest that, although the primary source of gene expression variation is tissue, HFD feeding significantly alters the gene expression profiles of APCs.
## I identified 1374 upregulated and 1291 downregulated genes in VWAT_HFD compared to SWAT_HFD, with an adjusted p-value < 0.05 and a log2 fold change> 0.06 (representing at least a 1.5 times difference). GO enrichment analysis revealed that VWAT_HFD enriched genes in pathways related to nuclear divisions and sister chromatid segregation, while reducing those associated with immune cells. GSEA analysis additionally indicated that VWAT_HFD activated genes involved in the mitotic spindle assembly checkpoint and regulation of chromosome segregation, but suppressed pathways linked to immune response regulation.
## In the original article, the authors revealed that HFD feeding increases the proliferation of APC in VWAT but not in SWAT. My analyses also showed that VWAT_HFD increased the expression of genes associated with cell divisions compared to SWAT-HFD. While we need to consider the tissue-specific gene expression patterns, my analysis replicates their findings. Interestingly, VWAT_HFD decreased the expression of immune system-related genes.
## The APC regulates immune responses and influences the expansion and function of adipose tissue. Our results suggest that consumption of a high-fat diet affects immune function in a tissue-specific manner. Further studies focusing on this axis are needed.
```
